# Relative importance of region, seasonality and weed management practices effects on the functional structure of weed communities in French vineyards

**DOI:** 10.1101/2021.10.26.465884

**Authors:** Marie-Charlotte Bopp, Elena Kazakou, Aurélie Metay, Guillaume Fried

**Affiliations:** CEFE, Univ Montpellier, CNRS, EPHE, IRD, Univ Paul Valéry Montpellier 3, Institut Agro, Campus CNRS/CEFE, 1919 route de Mende 34293, Montpellier, France; UMR ABSys, Institut Agro, Inra, Cirad, CIHEAM-IAMM, Univ Montpellier, 2 Place Pierre Viala, 34060 Montpellier, France; Anses, Laboratoire de la Santé des Végétaux, Unité Entomologie et Plantes invasives, 34988 Montferrier-sur-Lez, France

**Author notes:** **Corresponding author:** Marie-Charlotte Bopp, Centre d’Ecologie Fonctionnelle et Evolutive UMR 5175, Campus CNRS/CEFE, 1919 route de Mende 34293 MONTPELLIER CEDEX 5.

**Keywords:** trait-based approach, French wine-growing regions, seasonality, tillage, chemical weeding, mowing

## Abstract

Winegrowers have diversified their weed management practices over the last two decades changing the structure and the composition of weed communities. Complementary to taxonomic studies, trait-based approaches are promising ways for a better understanding of weed communities’ responses to environmental and agronomic filters. In the present study, the impact of climate, soil characteristics, seasons and weed management practices (chemical weeding, tillage and mowing) were assessed on weed communities in three French wine-growing regions (Champagne, Languedoc and Rhône valley). These agro-environmental gradients structuring weed communities according to their combination of traits were highlighted using multivariate analysis (RLQ). The impacts of these filters on Community Weighted Means (CWM) and the Community Weighted Variance (CWV) of weed communities were analysed using mixed and null modelling. Our results showed that spatio-temporal and weed management practices variables explained from 13% to 48% of the total variance of CWM (specific leaf area, maximum height, seed mass, flowering onset and duration and lateral spread). Region, seasonality and management practices explained 53%, 28% and 19% of CWM marginal variance, respectively. Weed management impacted CWM and CWV through two main gradients: (i) a soil disturbance gradient with high mechanical disturbance of soil in tilled plots and low mechanical disturbance in chemically weeded plots and (ii) a vegetation cover gradient with high vegetation abundance in mowed plots compared to more bare soils in tilled and chemically weeded plots. Chemically weeded communities showed trait values of ruderal strategies (low seed mass, small-stature) in Languedoc. Mowed plots were associated with more competitive strategies (higher seed mass, higher stature and lower SLA) in Languedoc. Tillage favoured communities with high seed mass that increases the viability of buried seeds and high lateral spread values associated to the ability to resprout after tillage in Languedoc and Champagne. This study demonstrated that trait-based approach can be successfully applied to perennial cropping systems such as vineyards, in order to understand community assembly to better guide weed management practices.

## 1. Introduction

Weed communities have an important role in maintaining biodiversity in agroecosystems, potentially delivering multiple ecosystem services as long as their negative impact on crops are limited (Gaba et al., 2015; Paiola et al., 2020; Storkey and Westbury, 2007; Winter et al., 2018). Understanding how weed communities respond to abiotic, biotic and anthropogenic factors is essential to better guide weed management practices and thus to increase their ecosystem services (e.g. climate regulation through carbon stockage, nitrogen supply) and decrease their ecosystem disservices (e.g. competition for soil water) (Mahaut et al., 2020).

In vineyards, winegrowers have diversified their weed management practices over the last two decades (Fernández-Mena et al., 2021; Novara et al., 2018; Simonovici, 2019). Chemical weeding, especially in inter-row, is less applied in favour of superficial tillage and mowing. These alternative practices have impacted the structure of weed communities (Fried et al., 2019; Gago et al., 2007; Steenwerth et al., 2016). For instance, the combination of tillage and mowing of inter-rows was significantly associated with higher richness and abundance than tillage or chemical weeding (Fried et al., 2019).

In addition to the taxonomic characterization of weed communities, trait-based approach can be used to explain the response of weed communities to environmental filters and weed management practices (Kazakou et al., 2016). Traits are any morphological, physiological or phenological features measurable at the individual level, from the cell to the whole-organism level (Violle et al., 2007). As other environmental drivers (e.g. climate, soil characteristics, seasonality), weed management practices select weed species within communities according to their trait values, so-called “response trait” (Damour et al., 2018; Kazakou et al., 2016; Lavorel and Garnier, 2002). Filtering processes can shape functional structure of weed communities in two major dimensions. Firstly, the mean trait values of communities reflect the major weed strategy to adapt to local conditions (e.g. early flowering onset to avoid perturbations). Secondly, the range of possible trait values expresses the dissimilarity of weed strategies within a community (e.g. wide range of flowering onset within a community might increase the probability that some species flower in a disturbed environment). Reduced or wide range of possible trait values lead respectively to convergent or divergent distributions (Bernard-Verdier et al., 2012; Perronne et al., 2017).

So far, trait-weed management practices relationships have been mostly explored in annual cropping systems (Alarcón Víllora et al., 2019; Armengot et al., 2016; Fried et al., 2012; Hernández Plaza et al., 2015; Smith, 2006; Storkey et al., 2010; Trichard et al., 2013) while few studies have investigated it in perennial crops as vineyards (Fiera et al., 2020; Hall et al., 2020; MacLaren et al., 2019; Mainardis et al., 2020). In vineyards, tillage, chemical weeding and mowing are the most frequent weed management practices applied in France (Simonovici, 2019). Tillage and chemical weeding can be considered as high disturbances as they destroy pre-existing living plant biomass (Gaba et al., 2014; Grime, 1979). Highly disturbed soil can result in convergent trait values distribution within the communities and favour trait values associated to ruderal weed communities (species with short stature, small seeds and high specific leaf area) (Grime, 2006; Kazakou et al., 2016). In contrast, mowing only partially destroys weed biomass. It may correspond to an intermediate disturbance (Grime, 2006), leading to a more divergent trait distribution (i.e. increased range of possible trait values) and to more competitive weed communities as vegetation cover is expected to be higher (species with large stature and high seed mass) (Kazakou et al., 2016; Mainardis et al., 2020).

Moreover, other abiotic filters such as climate, soil characteristics or seasonality can impact strongly the functional structure of weed communities (Keddy, 1992). Firstly, climate (e.g. temperature, precipitation) affects functional metrics at the community level (Alarcón Víllora et al., 2019; Hall et al., 2020). For instance, Alarcón Víllora et al. (2019) found that climatic inter-annual fluctuations drove the functional structure of weed communities more than management practices in cereal-legume rotation. Secondly, seasonality was one the main factor explaining weed community composition in annual crop fields (Fried et al., 2008; Hallgren et al., 1999; Lososová et al., 2004). However, few studies have explored the relative importance of those abiotic filters compared to weed management practices on functional structure of weeds in vineyards. Indeed, most studies have been made at the experimental level (except (Hall et al., 2020)) in fixed abiotic conditions without considering the effect of pedoclimatic variations.

In vineyards, some recent trait-based studies have considered functional diversity using various metrics (e.g. richness, evenness, divergence, dispersion) (Fiera et al., 2020; Hall et al., 2020; Mainardis et al., 2020). However, to the best of our knowledge, none of these studies have tested the filtering effect of weed management practices on variations in traits values of weed communities by using null models. Null model approach are largely used in community ecology to detect signatures of niche-based mechanisms (Perronne et al., 2017) and could be applied to managed weed communities in vineyards.

In this study, the relative importance of wine-growing regions (covering a wide range of climate and soil characteristics), seasonality and weed management practices on the functional structure of weed communities was assessed using Fried et al. (2019) large data set of 46 vineyards (so called Biovigilance network) from three wine-growing French regions (Champagne, Languedoc and Rhône valley). The general objective of our study was to test whether changes in weed species composition in vineyards caused by environmental and anthropogenic filters shown in Fried et al. (2019) would also lead to changes in functional structure. First, we highlighted the main agro-environmental gradients structuring weed communities according to their combination of traits, using multivariate analysis (RLQ). Then, two different aspects of the functional structure of the weed communities were assessed using trait values from databases: Community Weighted Means (CWM) which is the mean value of traits of weed communities, and Community Weighted Variance (CWV) which is the variability of these trait values within the community. We used mixed linear models to test the effects of the explanatory variables on the CWM of weeds communities. Secondly, we evaluated the seasonality and weed management practices effects on CWM within each region. Thirdly, we tested if CWV were significantly impacted by weed management practices and seasonality using a null model approach to disentangle the effect of functional variance from the effect only due to species richness (Perronne et al., 2017). We expected that seasonality and region would explain more CWM variability than weed management practices. We hypothesized that tillage and chemical weeding would restrict the range of possible trait values within weed communities and favour a ruderal strategy. On the contrary, we hypothesized that mowing would increase functional diversity and favour a more competitive strategy. Moreover, we hypothesized that intraspecific variation was lower than interspecific variation (species robustness assumption) (Garnier et al., 2001; Kazakou et al., 2014).

## 2. Materials and methods

### 2.1. Climate, soil characteristics and weed management practices

Weed surveys were performed in 46 vineyards from 2006 to 2012 in three main wine production regions in France (so called “Biovigilance network”): i) Champagne, northeast France (10 plots) ii) Beaujolais and northern Rhône valley, central east France (18 plots), and iii) Languedoc, central south France (18 plots). The climate of Champagne is continental with oceanic influences (Supplementary table 1). The mean annual temperature of Champagne is 10.1°C with 657 mm annual rainfall in the surveyed plots (Supplementary table 1). The climate of Rhône valley is semi-continental with a mean annual temperature of 11.4°C and 776 mm annual rainfall in the surveyed plots. The climate of Languedoc is Mediterranean with a mean annual temperature of 14.1 °C and 686 mm annual rainfall in the surveyed plots.

The soils of the Champagne vineyard plots are silty (45.7 %) with a neutral pH (pH of 7.1) with low bulk density (fine earth) mean value (1387.3 kg/m^3^) (Supplementary table 1). Rhône vineyards soils are characterized by the highest soil organic carbon content (19.7 %) with a slightly acidic pH (6.7). Languedoc plots soils are basic (pH of 7.5), have a high bulk density (1528 kg/m^3^) and are clayey (27% of clay). Because of a strong correlation between regions and pedoclimate variables, we have chosen to keep only the “region” variable, assuming that it largely represents the pedo-climatic differences.

Three different weed management practices were applied on rows and inter-rows in these vineyards: chemical weeding, tillage and mowing. As mowing on rows was only exceptionally applied in our dataset (applied in two plots in Rhône, representing 7 floristic surveys), we decided not to consider this variable. At the global dataset scale, chemical weeding concerned one third of the inter-rows and 90% of the rows. Farmers of the vineyard network used pre-emergence and post-emergence herbicides. Active ingredients of post-emergence herbicide were mostly glyphosate. Pre-emergence herbicide was mostly constituted by oryzalin. Tillage was applied on one third of the inter-rows and 17% of rows. Tillage was mostly superficial (mean of 12 cm and range from 5 cm to more than 20 cm). Mowing concerned one third of inter-rows.

Weed management practices differed according to wine-growing regions. In Languedoc, tillage was more frequent (70% of inter-rows practices and 27% of rows practices) than chemical weeding (26% of inter-rows practices and 85% of rows practices) and mowing (13% of inter-rows practices) (Table 1). On the contrary, tillage was less frequent in Rhône plots (7% of inter-rows practices and 9% of rows practices). Most of the rows were chemically weeded in this region (95%) and inter-rows were chemically weeded (45%) and mowed (52%). Inter-rows located in Champagne were mostly mowed (63%) followed by chemical weeding (48%). 84% of rows in Champagne were chemically weeded and 17% were tilled.

**Table 1.** Characteristics of weed management practices of rows and inter-rows in Champagne, Languedoc and Rhône. Combination of different management practices can be applied on rows or inter-rows so total percentage per region are not equal to 100%.

| Location of<br>weed<br>management<br>practices | Weed<br>management<br>practices | Abbreviations | Champagne<br>(10 plots) | Languedoc<br>(18 plots) | Rhône<br>(18 plots) |
| --- | --- | --- | --- | --- | --- |
| **Inter-rows** | Chemical<br>weeding | Chem.IR | 48% | 26% | 45% |
|  | Mowing | Mow.IR | 63% | 13% | 52% |
|  | Tillage | Till.IR | 28% | 70% | 7% |
| **Rows** | Chemical<br>weeding | Chem.R | 84% | 85% | 95% |
|  | Tillage | Till.R | 17% | 27% | 9% |

### 3.2. Floristic surveys

From 2006 to 2012, floristic surveys were performed in late winter to early spring (January to April), summer (May to July) and late summer to early autumn (August to October). Two temporal variables were considered in this study: the year of floristic survey and the number of days between the 1^st^ January of the same year and the day of the floristic survey, which is considered as an indicator of the seasonality. In each vineyard plot, plant species were surveyed over an area of 2000m^2^ (see Fried et al. (2019) for more details). To estimate species abundance, we used five abundance classes developed in Barralis (1976) and transformed into a quantitative scaling as followed: “1”, 0.5 individual/m^2^; “2”, 1.5 individuals/m^2^; “3”, 11.5 individuals/m^2^; “4”, 35.5 individuals/m^2^; “5”, 75 individuals/m^2^. A list of species and distinct abundance scores were noted for rows and inter-rows. However, in this study, we focused at the plot-scale flora resulting from the combination of row and inter-row practices (following MacLaren et al. (2019)). Therefore, plant community composition was estimated from the whole 2000 m^2^ surveyed including both the row and the inter-row (hereafter vineyard plot scale) taking the maximum abundance score for species occurring in both areas. In total, 270 surveys were recorded at the vineyard plot scale (46 in Champagne, 102 in Languedoc and 122 in Rhône).

### 2.2. Traits data

Six plant traits were selected to capture plant responses to environmental variations and weed management practices. Three traits of the Leaf-Height-Seed (LHS) strategy scheme were selected (Westoby, 1998): (a) specific leaf area (SLA) which is the light-catching area deployed per unit of previously photosynthesized dry mass, is related to the speed of resources acquisition (Wright et al., 2004), (b) maximum height which expresses the possible amount of growth in an undisturbed environment and which is related to light and nutrient acquisition (Westoby et al., 2002), (c) seed mass which represents the “colonisation-competition” trade-off (Moles and Westoby, 2006) illustrating two strategies: “producing a large number of small seeds, each with low establishment ability and high colonizing capacity” and “producing fewer, larger seeds, each with a higher chance of successful establishment” (Westoby et al., 2002). Three other traits related to persistence and regeneration in disturbed habitats were selected: (d) flowering onset, (e) flowering duration and (f) lateral spread ability. Lateral spread is a qualitative trait which represents species abilities to develop horizontally (species with rhizomes or forming tussocks); it is rated as followed: “1”, therophytes; “2”, perennials with compact unbranched rhizomes or forming small tussocks (less than 100 mm in diameter); “3”, perennials with rhizomatous system or tussocks reaching from 100 to 250 mm; “4”, perennials reaching diameter of 251 to 1000 mm.

The traits values were extracted from different databases: the LEDA Traitbase for SLA (Kleyer et al., 2008), Flora Gallica for maximum height (Tison and De Foucault, 2014), the Seed Information Database (SID) for seed mass (Royal Botanic Gardens Kew, 2021), Baseflor for flowering onset and duration (Julve, 1998) and lateral spread from Hodgson et al. (1995) supplemented by expert opinion (G. Fried, pers. com.).

We calculated the community weighted means (CWM) (Garnier et al., 2004) and the Community Weighted Variances (CWV) (Sonnier et al., 2010) of each trait at the vineyard plot scale using the following equations:

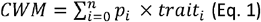

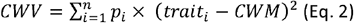

Where *p_i_* is the proportion of total abundance of the species *i* within a community, *trait_i_* is the value of trait of the species *i* and *n* is the total number of species within the community. CWM as the weighted average trait values of the community, expresses the most probable trait value of an individual randomly picked up within a community (Garnier et al., 2004). CWV expresses the variability of the trait values around the average value within the community (Sonnier et al., 2010).

### 3.3. Data analyses

#### 3.3.1. RLQ and fourth-corner analysis

To characterize the covariation of the functional structure of weed communities with management practices and spatio-temporal variables (i.e. region, seasonality and the year of floristic survey), we combined an RLQ analysis and a fourth-corner approach using Dray et al. (2014) framework. RLQ analysis investigates co-inertia between 3 types of data: i) region, year and season of floristic surveys (i.e. the number of days since the beginning of the year) and soil management variables (R table), ii) floristic composition (L table), iii) species trait attributes (Q table). Species density were square-root transformed. Firstly, correspondence analysis (CA) was applied to the table L. Then, we performed a Hill and Smith analysis on the R and Q tables using respectively the row and the column scores of the CA. Finally, the RLQ performed two co-inertia analyses on the R-L and L-Q tables. A Monte Carlo permutation (n=999) test was used to test the global significance of the relationship between the agro-environmental table R and the trait table Q. Based on the results of the RLQ analysis, a fourth-corner analysis was performed to test the significance of the relationship between traits and agro-environmental variables. At this step, we tested the associations between individual traits and environmental gradients obtained from RLQ scores, and between individual environmental variables and trait syndromes obtained from RLQ scores. We used a permutation model (n=49 999) to test the null hypothesis that species are distributed independently of their traits values and their preferences for agro-environmental conditions in the vineyard plots (Dray and Legendre, 2008). Adjusted p-value were used rather than p-value to limit the overall error rate of multiple testing. Multivariate analyses such as RLQ give a good idea of the main agro-environmental gradients. To further understand the effect of each agro-environmental variable on each trait, we analysed the variations in CWM and CWV.

#### 3.3.2. Mixed linear models of CWM

To evaluate the relative importance of region, temporal variables and weed management practices effects on CWM, we constructed mixed linear models for each CWM (“lmer” function of lme4 package (Bates et al., 2015)). We defined two random effects in each model: the vineyard plot identity and the year of floristic survey. Seed mass, lateral spread and flowering duration were logged to validate hypotheses of linear models. Model selection was performed using a backward step selection procedure. We calculated the explained variance of each covariate as the percentage of variance additionally explained when each variable was added one by one to the model.

#### 3.3.3. Covariations between CWM and weed management practices and temporal variable gradients

As region had a major effect on CWM, we investigated the weed management practices variables, the seasonality and the year of survey effects on CWM within each region. To characterize the gradient of weed management practices and temporal variation of floristics surveys, we performed a Principal Component Analysis (PCA). Then, we tested the correlations between CWM and the PCA scores of the sites on the first two axes representing gradients of management practices (Spearman’s rank correlation). We corrected p-values from multivariate testing using Bonferroni corrections.

#### 3.3.4. Null modelling and covariations between effect sizes of CWV and weed management practices and temporal variable gradients

To test whether CWV values were not randomly distributed along the weed management practices gradient, we first used a null model approach. We constructed a “population-based fixed-zero per sites” null model to test the following null hypothesis: abundance is randomly distributed within plots with respect to trait values. We shuffled species x site matrix for the observed species, while keeping species x trait matrix unchanged, breaking the link between abundance and trait values (Bernard-Verdier et al., 2012; Perronne et al., 2017). Thus, the richness, the list of the observed species and the abundance distribution within a plot remained unchanged. This randomization type allows to disentangle the effects of environmental and agronomic drivers on functional diversity from effects simply related to the richness of communities. For each plot, we calculated an effect size (ES) as the quantile of the null distribution in which the observed value is found (Bernard-Verdier et al., 2012; Kelt et al., 1995) (Eq. 3).

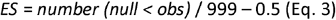

The effect sizes measure the strength of convergence and divergence (Botta-Dukát, 2018). ES values close to zero indicate that observed CWV values are not different from random CWV values. In contrast, high and low ES values quantify respectively strong divergent and convergent functional structure of weed communities. In order to detect general pattern of community structure regardless of the spatio-temporal and agronomic characteristics, we tested if ES was overall different from 0 using a two-tailed Wilcoxon signed-ranks test (Bernard-Verdier et al., 2012; Jung et al., 2010). To investigate the impact of the weed management practices gradient on CWV, we used the same procedure as for CWM. Within each wine-growing region, we tested the significance of correlations between effect sizes of CWV and the coordinates of the plots on the first two axes of the PCA, using Spearman’s rank correlation. All the statistical analyses were performed with R (3.6.2 version). All multivariate analyses (RLQ, PCA) were conducted using ade4 package (Chessel et al., 2004).

## 3. Results

### 3.1. Covariation of the functional structure of weed communities across management practices and spatio-temporal variables

The first two axes of the RLQ explained 95% of inertia (81 % explained by the first axis and 14% explained by the second axis) (Figure 1). The co-structure between R and Q was significant (Monte-Carlo test, P < 0.001) demonstrating the global significance of the relationships between species traits and agro-environmental variables (region, seasonality, year of floristic survey and weed management practices as specified in Table 1). According to the fourth-corner analysis combined with the RLQ analysis, all the spatio-temporal and agronomic variables except Rhône region were correlated to the first axis of the RLQ, which described most of the variability (Figure 2). The first RLQ axis opposed spring surveys and the later survey years, located in Champagne region with chemical weeding on the rows and either chemical weeding or mowing of inter-rows to autumn surveys and the first years of surveys, located in Languedoc region with tilling practices in both rows and inter-rows (Figure 1a, 2a).

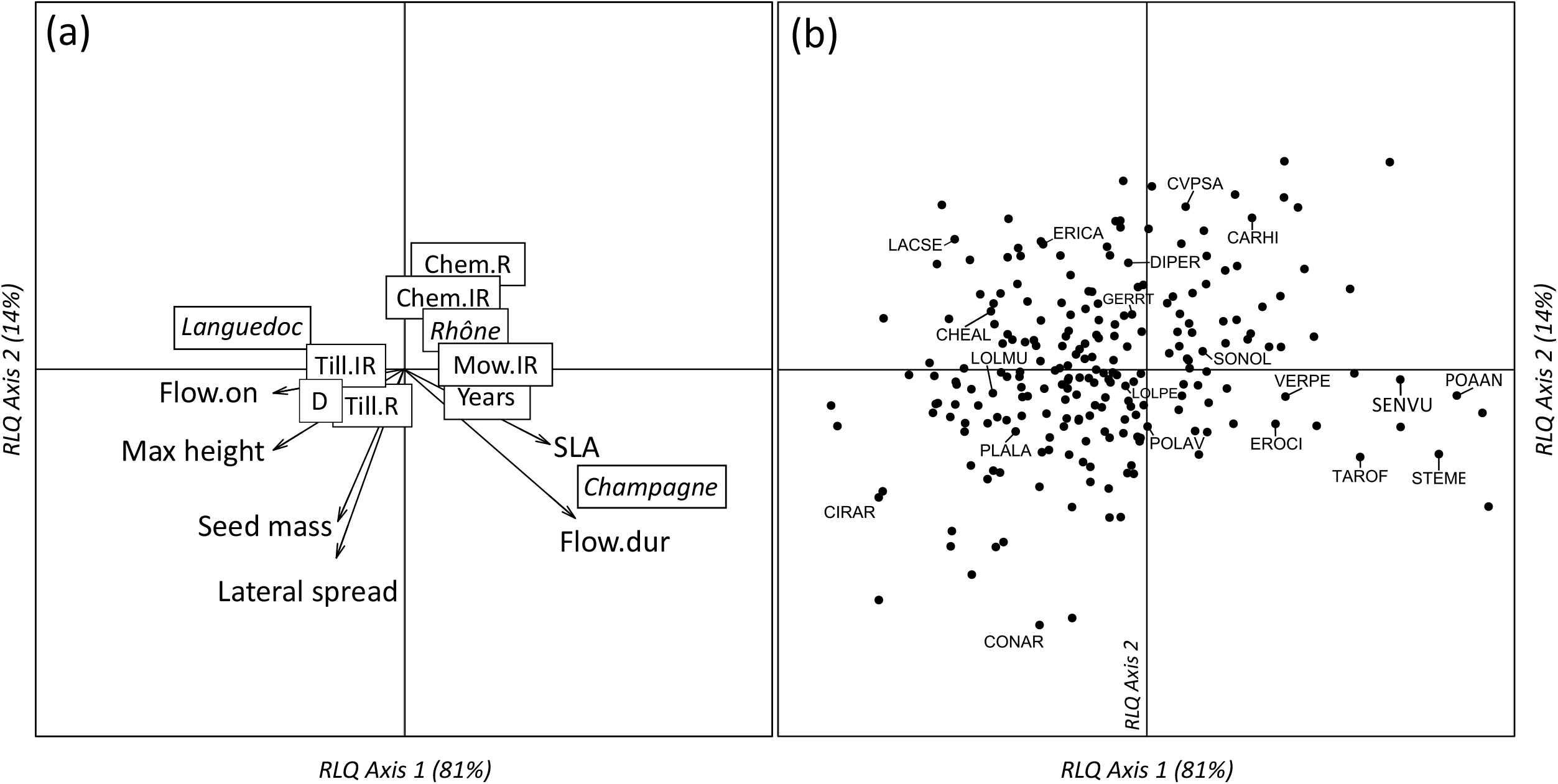

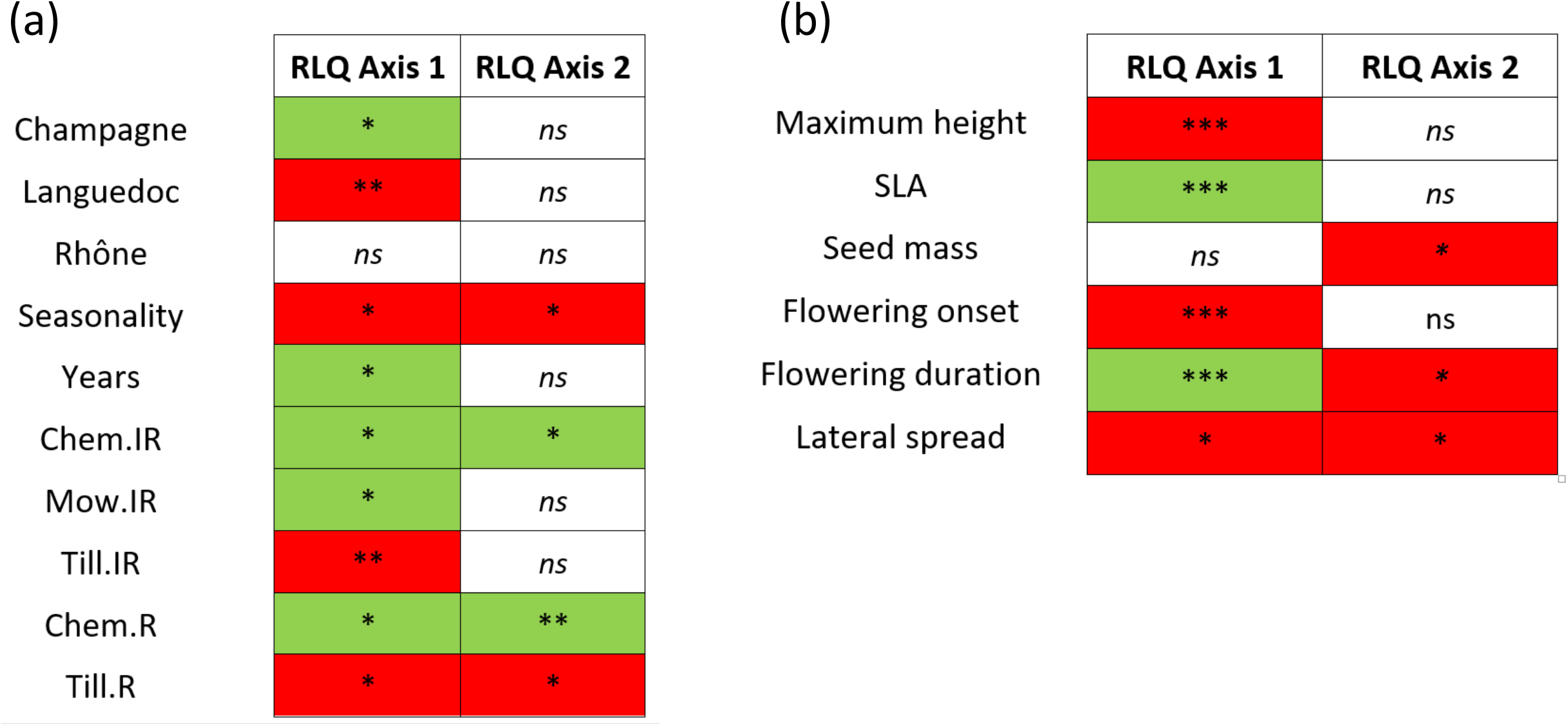

Species with short stature, small-sized seeds, high SLA and long flowering time (e.g. *Stellaria media* which has a flowering duration of 12 months and a SLA of 42.09 m^2^/kg) were associated with the Champagne region and spring surveys on the positive side of the first axis, favoured by chemical weeding on the rows and the inter-rows and mowing on the inter-rows (Figure 1a, 2a, 2b). On the opposite, species with high stature, large seeds, late flowering and high lateral spread abilities (e.g. *Cirsium arvense* which can reach 1.5 m high, with long horizontal roots favouring lateral spread and with first flowers appearing in July) were associated with the Languedoc region, autumn survey on the negative side of the first axis and favoured by tillage (Figure 1b). The second RLQ axis discriminated chemical weeding of rows and inter-rows to row tilling (Figure 1a, 2a). Species with high seed mass, long flowering duration and high lateral spread abilities (e.g. *Convolvulus arvensis* with rhizomes which favour lateral spreading and flowering duration of 6 months) were associated to tillage on the rows on the negative side of the second axis (Figure 1b, 2a, 2b). On the opposite side, species with small-sized seeds, short flowering duration and low lateral spread abilities (e.g. *Crepis sancta* with less than 1 mg seed weight and 3 months of flowering duration) were associated to chemical weeding of the rows and of the inter-rows (Figure 1b).

### 3.2. Relative importance of weed management practices and spatio-temporal variables explaining weed community’s functional response

Spatio-temporal and weed management practices variables explained from 13% to 48% of the total variance of CWM of the different traits (Figure 3). Overall, region explained most of CWM marginal variance (53%), followed by seasonality (28%) and some weed management practices variables (19%).

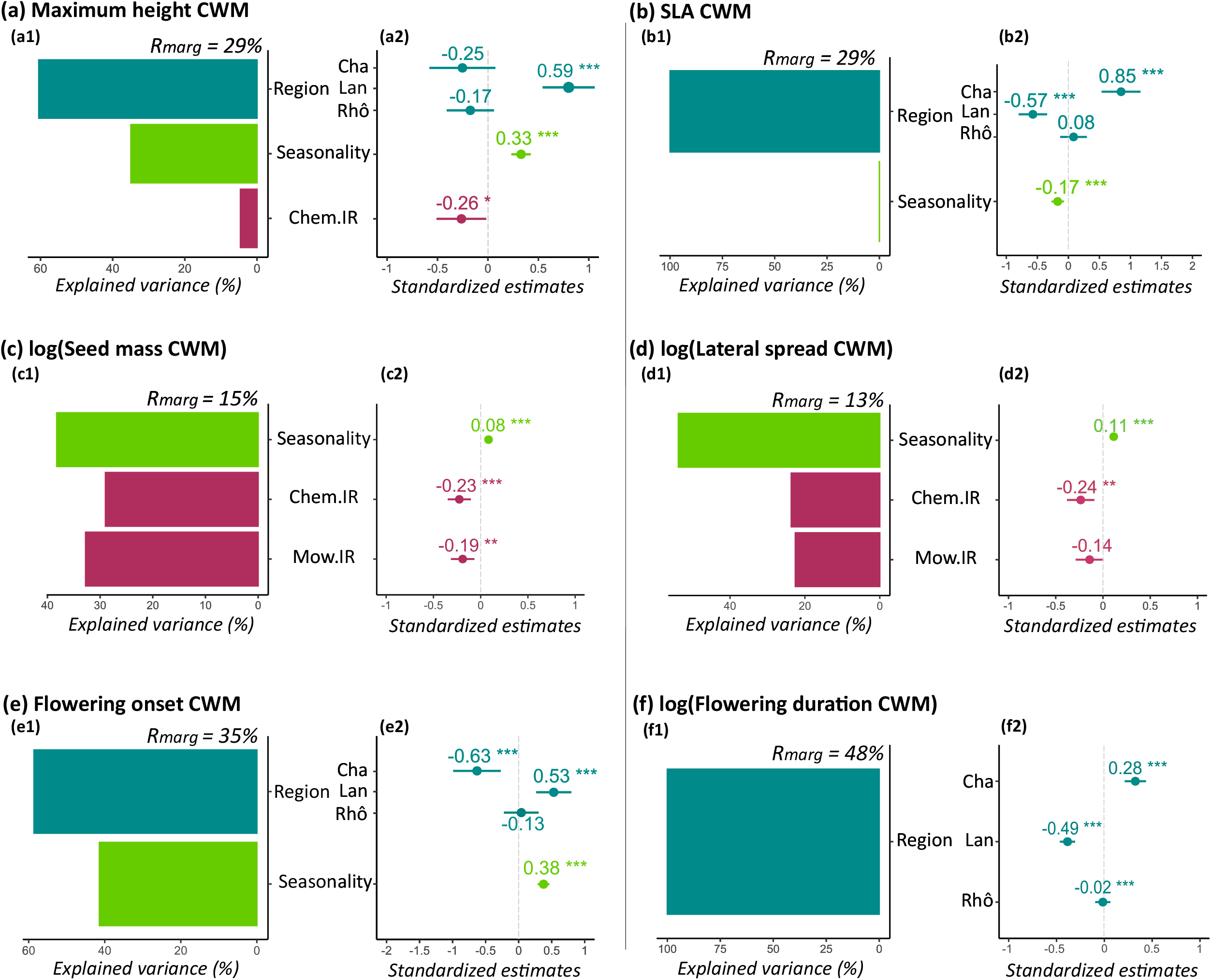

The region of floristic surveys explained large part of variance of maximum height (60%, Figure 3a1), SLA (99%, Figure 3b1), flowering onset (59%, Figure 3e1) and flowering duration (100%, Figure 3f1). This effect was not significant for seed mass and lateral spread. Weed communities from Champagne region had high SLA (estimate: 0.85, Figure 3b2), early flowering onset (estimate: −0.63, Figure 3e2) and long flowering duration (estimate: 0.28, Figure 3f2). Weed communities from Languedoc had low SLA (estimate: −0.57, Figure 3b2), late flowering onset (estimate: 0.53, Figure 3e2), short flowering duration (estimate: −0.49, Figure 3f2) and high stature (estimate: 0.59, Figure 3a2). The estimate of flowering duration in Rhône was almost null (−0.02) but the effect was significant showing that flowering duration was medium in that region and different from Champagne with short flowering duration and Languedoc with long flowering duration (Figure 3f2).

Seasonality was the most frequently selected effect in CWM models. It explained significant proportions of variance of CWM of maximum height (35%, Figure 3a1), seed mass (38%, Figure 3c1), lateral spread (54%, Figure 3d1), flowering onset (41%, Figure 3e1) but low variance of SLA (<1%, Figure 3b1). Seasonality did not significantly impact flowering duration. The communities of autumn floristic surveys had high stature (estimate: 0.33, Figure 3a2), high seed mass (estimate: 0.08, Figure 3c2), high lateral spread abilities (estimate: 0.11, Figure 3d2) late flowering onset (estimate: 0.38, Figure 3e2) and low SLA (estimate: −0.17, Figure 3b2).

Mowing and chemical weeding variables were selected in the mixed linear models. Chemical applications on inter-rows explained large proportions of variance of seed mass (29% of explained variance, Figure 3c1), lateral spread (24% of explained variance, Figure 3d1) and fewer variance of maximum height (5% of explained variance, Figure 3a1). Mowing of inter-rows explained 33% of seed mass variance (Figure 3c1) and 22% of lateral spread variance (Figure 3d1). Chemically weeded communities on inter-rows had low stature (estimate: −0.26, Figure 3a2), low seed mass (estimate: −0.23, Figure 3c2) and low lateral spread abilities (estimate: −0.24, Figure 3d2). Mowed weed communities in inter-rows showed low seed mass (estimate: −0.19, Figure 3c2) and low lateral spread tendency (estimate: −0.14, Figure 3d2). Tillage of rows and inter-rows had no direct effects on CWM of weed species and in general, the management of the rows did not impact significantly CWM of weed communities.

The plot random effect described significant proportions of total variance (35% of lateral spread abilities, 20% of seed mass, 13% of flowering duration, 10% of SLA, 9% of maximum height, 8% of flowering onset). The random effect of the year of the floristic survey was only selected in the flowering onset CWM model and represented 10% of the total variance of this CWM.

### 3.3. Functional response of weed communities to weed management practices within each region

#### 3.3.1. Community Weighted Means (CWM) response to weed management practices within each region

In order to disentangle the effect of region from the effects of the other variables, we explored weed functional responses to weed management practices, seasonality and years of surveys within each region. Figure 4 displays the gradients of these variables, excluding the region effect. The first two axes represented 54% of total variance. They described mostly weed management practices gradients (Supplementary table 2). Seasonality and years of survey variables contributed poorly to total inertia of these axes (7% of explained variance for the first two axes). The first axis explaining 31% of variance opposed tilled rows and inter-rows (positive coordinates) and chemically weeded rows (negative coordinates). It represented the soil disturbance gradient from tilled soils with high below-ground mechanical disturbances to chemically weeded soils with low below-ground mechanical disturbance. The second axis explaining 23% of variance opposed mostly mowed inter-rows (negative coordinates) to combinations of tilled and chemical weeded inter-rows (positive coordinates). It represented the vegetation cover gradient with high vegetation cover in mowed inter-rows and low vegetation cover in tilled and chemically weeded inter-rows.

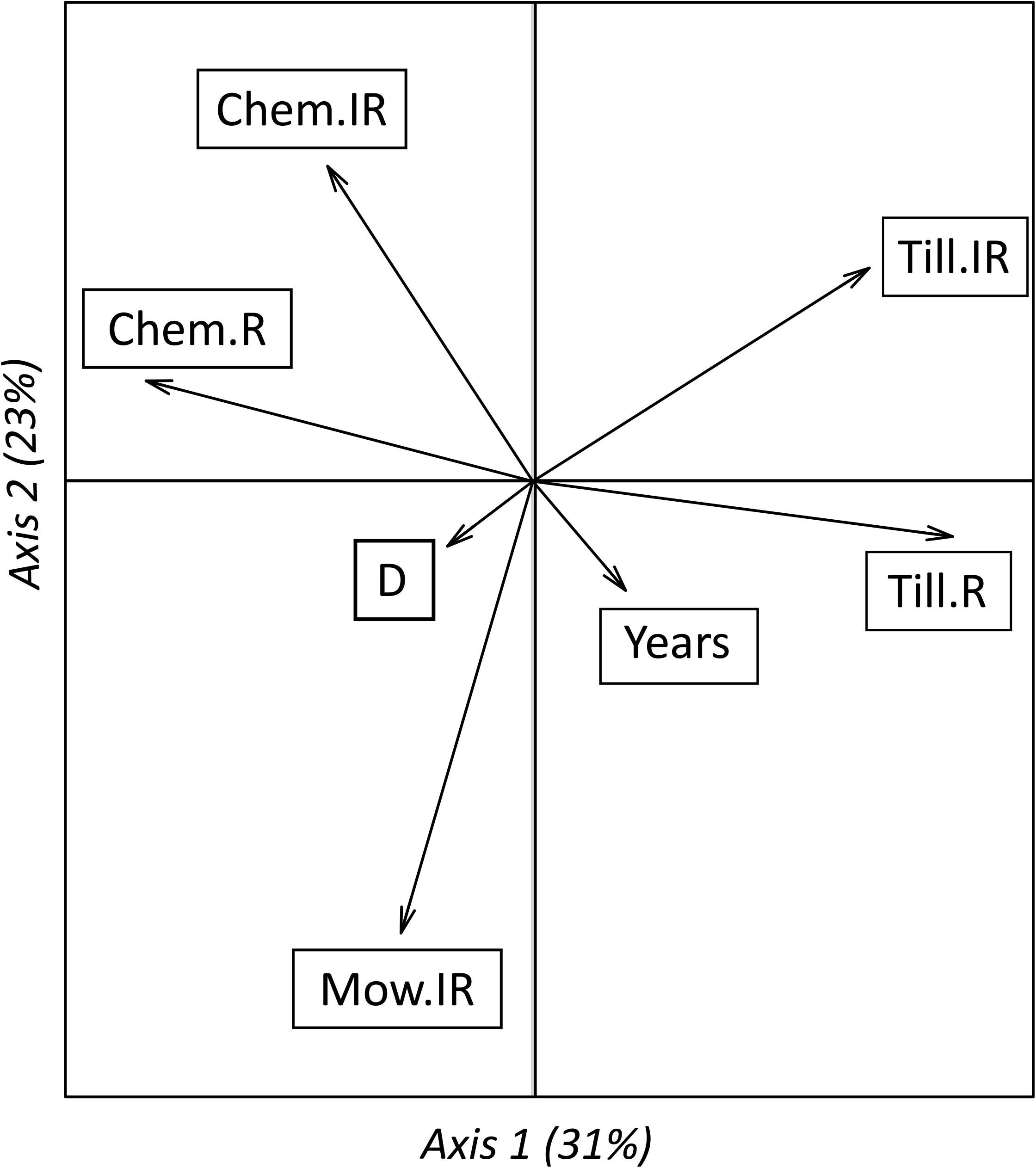

Table 2 shows the significance and values of coefficients of correlation between CWM within each region and the first two axes of the PCA performed on weed management practices, season and years of survey variables (Figure 4). Management practices effect on CWM differed according to the wine-growing regions (for means and standard deviations of CWM per region, see Supplementary table 3).

**Table 2.** Coefficients of correlation between Community Weighted Means (CWM) and weed management gradients (two first PCA axes, see Figure 4) for each region. P-values were corrected from multiple comparisons testing using Bonferroni correction. The first PCA axis opposed chemical weeding of rows (Chem.R, negative coordinates,) and tillage of rows and inter-rows (Till.IR, Till.R, positive coordinates). The second PCA axis opposed mowing of inter-rows (Mow.IR, negative coordinates) to combination of tillage and chemical weeding of inter-rows (Chem.IR + Till.IR, positive coordinates). * *p* < 0.05; ** *p* < 0.01; *** *p* < 0.001; no asterisks, non-significant (*p* > 0.05). SLA, Specific Leaf Area; PCA, Principal Component Analysis.

| CWM | Languedoc |  | Champagne |  | Rhône |  |
| --- | --- | --- | --- | --- | --- | --- |
|  | 1<sup>st</sup> PCA axis | 2<sup>nd</sup> PCA axis | 1<sup>st</sup> PCA axis | 2<sup>nd</sup> PCA axis | 1<sup>st</sup> PCA axis | 2<sup>nd</sup> PCA axis |
|  | Chem.R → | Mow.IR → | Chem.R → | Mow.IR → | Chem.R → | Mow.IR → |
|  | Till.IR, Till.R | Chem.IR +<br>Till.IR | Till.IR, Till.R | Chem.IR +<br>Till.IR | Till.IR, Till.R | Chem.IR +<br>Till.IR |
| **Maximum height** | 0.06 | -0.29 ** | 0.41* | -0.2 | -0.25* | -0.05 |
| **SLA** | -0.12 | 0.31** | -0.17 | 0.26 | 0.09 | 0.06 |
| **Seed mass** | 0.26* | -0.4*** | 0.001 | 0.04 | -0.01 | -0.16 |
| **Lateral spread** | 0.53*** | -0.32** | 0.36* | -0.21 | -0.09 | -0.21 |
| **Flowering onset** | 0.01 | -0.24* | 0.07 | -0.34 | -0.33*** | 0.01 |
| **Flowering duration** | 0.05 | 0.16 | -0.01 | 0.23 | 0.24* | -0.22 |

In Languedoc, significantly higher CWM of seed mass and lateral spread abilities were found in tilled rows and inter-rows compared to chemically weeded rows (Table 2). Mowing was significantly associated with lower CWM for SLA and higher CWM for maximum height, seed mass, lateral spread abilities and flowering onset compared to chemically weeded inter-rows and to combined tillage and chemical weeding of inter-rows. In Champagne, tillage on rows and inter-rows was associated with higher lateral spread abilities as in Languedoc region and higher maximum height compared to chemically weeded rows (Table 2). Mowing was not significantly correlated with CWM variation. In Rhône, chemical weeding on rows was significantly associated with shorter flowering duration, higher stature and later flowering compared to tillage of rows and inter-rows (Table 2). As in Champagne, mowing was not linked with CWM variation.

#### 3.3.2. Community Weighted Variance (CWV) response to weed management practices within each region

Half of the CWV were significantly different from random expectations of null models (Supplementary figure 7, 8, 9, 10, 11, 12). More precisely, most of the CWV were lower than expectations demonstrating a convergent distribution and a restricted variance of traits values within weed communities (for means and standard deviations of CWV per region, see supplementary table 4).

In Champagne, lateral spread CWV were convergent while flowering onset and duration had divergent distributions (Supplementary figure 10). In Languedoc, SLA, lateral spread, flowering onset and seed mass had convergent distributions (Supplementary figure 7, 8). In Rhône, seed mass and lateral spread were convergent (Supplementary figure 11, 12). Four effect sizes out of 36 were significantly correlated to one axis of the PCA (Table 3) demonstrating different functional responses to filtering effect of weed management practices. In Languedoc, the effect size of lateral spread CWV was positively correlated with the first axis, showing that species located in chemically weeded rows communities had similar lateral spread abilities while species within tilled communities had dissimilar lateral spreading strategies (Table 3). In Champagne, the effect sizes of SLA CWV, flowering onset CWV and flowering duration CWV were positively correlated with the second axis (Table 3) demonstrating that combination of chemical weeding and tillage of inter-rows was associated with high variations of SLA, flowering onset and duration within weed communities.

**Table 3.** Coefficients of correlation between effect sizes of Community Weighted Variance (CWV) and weed management gradients (two first PCA axes, see Figure 4) for each region. P-values were corrected from multiple comparisons testing using the Bonferroni correction. The first PCA axis opposed chemical weeding of rows (Chem.R, negative coordinates,) and tillage of rows and inter-rows (Till.IR, Till.R, positive coordinates). The second PCA axis opposed mowing of inter-rows (Mow.IR, negative coordinates) to combination of tillage and chemical weeding of inter-rows (Chem.IR + Till.IR, positive coordinates. * *p* < 0.05; ** *p* < 0.01; *** *p* < 0.001; no asterisks, non-significant (*p* > 0.05). SLA, Specific Leaf Area; PCA, Principal Component Analysis.

| Effect sizes | Languedoc |  | Champagne |  | Rhône |  |
| --- | --- | --- | --- | --- | --- | --- |
| of CWV | 1<sup>st</sup> PCA axis | 2<sup>nd</sup> PCA axis | 1<sup>st</sup> PCA axis | 2<sup>nd</sup> PCA axis | 1<sup>st</sup> PCA axis | 2<sup>nd</sup> PCA axis |
|  | Chem.R → | Mow.IR → | Chem.R → | Mow.IR → | Chem.R → | Mow.IR → |
|  | Till.IR, Till.R | Chem.IR +<br>Till.IR | Till.IR,<br>Till.R | Chem.IR +<br>Till.IR | Till.IR,<br>Till.R | Chem.IR +<br>Till.IR |
| Maximum height | 0.02 | -0.01 | 0.03 | 0.08 | -0.1 | 0.07 |
| SLA | 0.09 | 0.11 | -0.26 | 0.4 * | -0.08 | 0.09 |
| Seed mass | -0.06 | -0.06 | -0.07 | 0.02 | -0.01 | 0.03 |
| Lateral spread | 0.45*** | -0.19 | 0.17 | 0.08 | -0.11 | -0.03 |
| Flowering onset | 0.1 | 0.01 | -0.12 | 0.38* | -0.19 | 0.03 |
| Flowering duration | 0.03 | -0.11 | -0.08 | 0.41* | -0.07 | -0.09 |

## 4. Discussion

Our study highlighted that the functional structure of vineyard weed communities responded strongly to agro-environmental factors with high variation of trait values across regions, seasons and management practices. In addition to average trait values, we explored the filtering effect of weed management practices through the analysis of community weighted variances. To our knowledge, this is the first study to investigate weed management practices effects on functional structure of weeds through null modelling of community weighted variances in vineyards. Null modelling has allowed us to test if functional variance (CWV) were restricted or increased by weed management, independently to weed management effect on community’s richness has shown in Fried et al. (2019). This method, widely applied on natural ecosystems, are still sparsely applied in agricultural contexts, especially in vineyards. We hypothesized that chemical and tillage would act as stronger filters compared to mowing, and we expected that communities under chemical and tillage would have convergent values of trait values with low variation. Half of the community weighted variances had a significantly different distribution than random distribution and most of the community weighted variances had convergent distribution. This demonstrates that regions, seasonality, years of survey and weed management practices impacted traits variabilities, mostly restricting the possible range of values of average trait values of weed communities. Moreover, it is likely that the assembly of species into communities in vineyards, which remains a disturbed environment, is also the result of neutral processes related to spatial colonization–extinction dynamics as shown in annual crop fields (Perronne et al., 2015).

### 4.1. Region and seasonality are the main drivers of the variation of community weighted means

Region was the main driver of traits related to resource acquisition (maximum height, SLA) and phenology (flowering onset and flowering duration) while season explained most of the variation of the regenerative traits (seed mass and lateral spread). According to our hypotheses, region and seasonality affected more communities functional structure than management practices.

Region effect encompasses different environmental effects, mostly climate and soil characteristics differences. In the literature, the edaphoclimatic effects depend on the scale of studies. Within a same wine-growing region in South Africa, MacLaren et al. (2019) found no significant effects of soil characteristics and climate on communities weighed means. In contrast, in several European countries, Hall et al. (2020) found that the country effect was the main factor affecting traits. Within the same country, our study showed that divergent edaphoclimatic conditions between wine-growing regions had major impacts on traits.

More precisely, region effect encompasses particularly the differences of climate between Languedoc, Rhône and Champagne and had contrasted effects on community weighted means of communities in Champagne (drier and colder climate) and communities in Languedoc (hotter climate). In Champagne, weed communities presented higher SLA values compared to Languedoc. This result is consistent with other studies at the individual scale showing that SLA was negatively correlated with mean annual temperature (Garnier et al., 2019; Moles et al., 2014; Wright et al., 2005, 2004) and positively with precipitations (Garnier et al., 2019). Indeed, plants with low SLA invest in the leaf structure to adapt to dry conditions (e.g. thick leaf blade to limit evapotranspiration; small and thick-walled cells) (Wright et al., 2005). In average, flowering onset were later in Languedoc compared to Champagne where most of the weed species flowered in spring. This could be explained by the bi-modality of the flowering period (Thompson, 2007) in Languedoc region. Two favourable periods are available to flower: (i) early spring before the summer drought and (ii) early autumn after the first rainfalls (Kummerow, 1983; Thuiller et al., 2004). Due to higher temperatures in autumn, more thermophilous species can flower and produce seeds during this season in the Mediterranean region (e.g. *Dittrichia viscosa, Heliotropium europaeum, Sorghum halepense, Tribulus terrestris*).

Moreover, soil characteristics are also encompassed in the region effect. In our dataset, vineyard plots located in Rhône had more fertile soil (high soil organic content) compared to vineyard plots of the other regions (Supplementary table 1). Fertile soils are usually associated with high acquisition rate, high SLA, high height at maturity and low seed mass (Garnier et al., 2016). However, only flowering duration was significantly different in Rhône while the other traits were not significantly different from the other regions.

In addition to different soils and climates, the region effect might also include some management practices and technical characteristics that vary according to wine-growing regions: the level of fertilization and amendments, cep density (higher density in Rhône than in Languedoc) or grape variety (Gamay and Syrah in Rhône, Cabernet Sauvignon and Grenache in Languedoc).

In addition to the regional effect, seasonality was the most frequently selected effect in reduced models explaining average traits variations. Interestingly, Fried et al. (2019) found that season was the main driver of the taxonomic composition of weed community in vineyards. Autumn surveyed weed communities had higher maximum height, lower SLA, heavier seeds, high lateral spread abilities and later flowering onset than the other survey dates. This can be explained by the changes of environmental conditions throughout the growing seasons (Wolkovich and Cleland, 2014): in late winter, disturbance is high in the vineyards (first weeding passages) with non-limiting weather conditions (regular rainfalls, increasing temperatures) selecting early-flowering species with rapid-growth and acquisitive strategies (high SLA, low stature, small seeds) while in summer, disturbing events as weeding passage are less frequent and weather conditions can lead to water stress or heat stress. Consequently, more stress-tolerant communities might develop during the summer drought with slow-growth and more conservative strategies (low SLA, high stature, large seeds).

### 4.2. Soil disturbance gradient: soil tillage versus chemical weeding

The first PCA axis of weed management practices and temporal variables (seasonality and years of floristic surveys) represented the soil disturbance gradient from tilled soils with high below-ground mechanical disturbances to chemically weeded soils with low below-ground mechanical disturbance.

According to our hypotheses, chemical weeding on rows was associated to traits values characterizing ruderal communities (low seed mass, short stature). In contrast, tillage favoured weed communities with high seed mass which is not a trait value expected for ruderal communities. Different mechanisms can explain these contrasted traits values between these two types of weed management. One possible explanation relates to the changes of seeds position in the soil which depend on the different weed management practices. Indeed, chemical weeding associated to no-till practices favours superficial position of seeds, whereas tillage commonly buries the seeds deeper into the soil. Unburied seeds are more likely to be eliminated by predation or impaired by desiccation. Therefore, under chemical weeding, producing a large number of small seeds might increase the probability that some of them survive. On the contrary, large seeds have a greater probability to germinate when they are buried (Benvenuti et al., 2001; Hernández Plaza et al., 2015; MacLaren et al., 2019). Under superficial tillage practices (as here in vineyards), large-seeded community have been found in several studies (Armengot et al., 2016; Hernández Plaza et al., 2015a; MacLaren et al., 2019).

Moreover, tillage is a soil disturbance (Gaba et al., 2014) that selects annual species with a seedbank but also species that can regenerate from fragments such as rhizome species (e.g. *Convolvulus arvensis* or *Cirsium arvense*) with high lateral spread abilities as highlighted here in two regions (Languedoc and Champagne). Tillage was also associated to high variability of lateral spread values within communities. Thus, tillage seemed to favour two different strategies (Supplementary figure 13): the main strategy is the ability to re-sprout using vegetative multiplication after tillage (high lateral spread communities). The second minor strategy is similar to chemically weeded communities: short life cycle communities with low lateral spread abilities.

### 4.3. Vegetation cover gradient: mowing versus tillage and chemical weeding

In contrast to Fried et al. (2019) findings on taxonomic composition, mowing has here been found to be a major driver of functional structure of Languedoc weed communities. Vegetation abundance in mowed inter-rows has been found to be much higher than in chemically weeded and/or tilled inter-rows (Fried et al., 2019) which expresses the differences between inter-rows with bare soils versus temporary or permanent vegetation cover (Hall et al., 2020).

Interestingly, the combination of chemical weeding and tillage on inter-rows favoured ruderal communities in Languedoc (short-stature, high SLA, low seed mass, low lateral spread abilities and early flowering) and were opposed to more competitive communities on mowed inter-rows. The sequential application of a belowground (soil tillage) and an aboveground (herbicides) action thus act as a severe disturbance for vegetation. It selected species with a rapid life-cycle that flower early to escape disturbances, have a high acquisitive strategy (high SLA), a low investment in vegetative part (low maximum height) and a massive production of small seeds to increase the probability that some survive (Grime, 1977; White and Pickett, 1985).

Mowing was associated with late flowering species communities compared to chemical weeding and tillage in Rhône. This could be explained by the timing of weed management practices. Indeed, farmers mowed usually later than the other weed management practices (early June on average for mowing, late April to May for chemical weeding and late May for tillage in Languedoc).

We expected that highly disturbed rows and inter-rows as tilled and chemically weeded rows would lead to a reduction in the range of trait values (convergent distributions) compared to mowed rows and inter-rows (divergent distributions) (Kazakou et al., 2016). In contrast with our expectations, combined tillage and chemical weeding were associated with high SLA variability, high flowering onset and duration community weighted variances while mowing was associated to more convergent distribution in Champagne. A possible explanation is that chemical weeding and tillage select contrasted strategies (e.g., geophytes with high lateral spread and therophytes) leading to higher variability of traits values when combining them (Supplementary figure 13).

Finally, this high timing variability of flowering with combined tillage and chemical weeding can be explained by the weed management practices timing variability. Indeed, mowing is usually applied in summer (June, July and August), in the Champagne plots. In contrast, chemical weeding and tillage are applied over a longer period of time (from March for chemical weeding to July for tillage). Thus, communities with contrasting flowering onset and duration might increase the probability that some species develop under chemical and tillage weeding.

### 4.4. Limits and perspectives

Trait-based approach is promising to better understand the functional shaping of weed communities by weed management practices. In our study, lateral spread ability of communities was certainly one of the major response traits. This unusual trait compared to more classical trait like SLA, height or seed mass allowed us to describe the assembly of the communities in a more mechanistic way. Other traits, absent from database, as the presence of epicuticular wax on leaves or seed coat thickness, might be response traits of interest to include in such studies where herbicide pressure is an important filter (Gaba et al., 2014). However, these specific traits are still lacking in trait databases. More effort should be done to expand trait databases (Kattge et al., 2020).

The use of pluriannual database of floristic survey as Biovigilance network is an asset to consider the variations between years due, for instance, to changing climatic conditions. Moreover, the wide geographic range of our study allowed us to analyse weed management practices and regions. At this large-scale of analysis, one drawback is that it is hardly possible to use measured trait values. We therefore used database traits values based on the assumption that the ranking of species according to their traits values is stable across environments (“stable species hierarchy”, Kazakou et al., 2014) as interspecific variability is higher than intraspecific variability. A recent study has shown that this hypothesis was largely valid in vineyard (Garcia et al., 2020).

Another point is that our results demonstrated that weed management practices explained 19 % of variations of functional structure of weed communities. More detailed variables describing management practices could help to a better understanding of plant responses, for instance considering disturbance types as frequency (e.g. number of applied management practices within a year) and intensity (e.g. herbicide dose or depth of tillage) (Gaba et al., 2014). For instance, timing of weed management practices has been shown to be relevant to explain functional responses of weed communities (Cordeau et al., 2017; Smith, 2006). Moreover, most of the theorical background on disturbance gradients have been done in grasslands and associated disturbances (e.g. grazing). More theorical frameworks are needed to better define weed management practices as disturbances and associated assumptions on trait variation (Gaba et al., 2014).

## 5. Conclusion

In this paper, we showed that trait-based approach can be successfully applied to better understand community assembly in an agricultural context. We showed that the changes in weed species composition caused by environmental and anthropogenic filters in vineyards also lead to changes in functional structure. Region, seasonality and weed management practices act as strong drivers of functional structure of weed communities. Weed management practices impacted both the mean traits values and their variance within weed communities. Chemically weeded communities showed trait values of ruderal strategies (low seed mass, small-stature). Mowed plots were associated with more competitive strategies (higher seed mass, higher stature and lower SLA). Tillage was associated with high seed mass that increase the viability of buried seeds and high lateral spread abilities values related to the capacity to resprout after tillage. Nowadays, mowing and tillage are more and more applied in vineyard (Simonovici, 2019). Our results showed that this soil management shift might favour more competitive communities. These weed communities might have different impact on agrosystem processes as nitrogen cycling or carbon sequestration through change of soil microbial composition (Karimi et al., 2020). Understanding the effect of weed communities on such processes is needed to adapt weed management practices and better drive ecosystem services and disservices (Damour et al., 2018; Garcia et al., 2018; Petit et al., 2018; Storkey et al., 2015).

## 6. Acknowledgement

This research was supported by Occitanie Region (Arrêté modificatif N° 19008795 / ALDOCT-000660 Subvention d’investissement, Allocations de recherche doctorales 2019) and the Office Français de la Biodiversité (ECOPHYTO II: Axe 2 – Action 8 and 9, N°SIREPA: 4148) as part of the SAVING project: Spatio-temporal dynamics of weed species communities in response to soil management practices in vineyards and consequences for grapevines: transition to zero glyphosate management. We would like to thank all winegrowers who provided management information and access to their farms. Thanks to the Biovigilance Flore network including all the people from SRAL and FREDON who performed the surveys, Nicolas André (FREDON Occitanie), Jacques Grosman (DGAL, SRAL Rhône-Alpes), and Olivier Pillon (SRAL Champagne) for data management at the regional level, and the Ministry of Agriculture for funding the monitoring. Warmful thanks to Margot Puiraveau who gathered the dataset in 2015. The study utilised data provided through the TRY initiative on plant traits (http://www.try-db.org). The TRY initiative and database is hosted, developed and maintained by J. Kattge and G. Beonisch (Max Planck Institute for Biogeochemistry, Jena, Germany). TRY is currently supported by DIVERSITAS/Future Earth and the German Centre for Integrative Biodiversity Research (iDiv) Halle-Jena-Leipzig. The authors have no conflicts of interest to declare.

