## Supplementary for "Relative importance of region, seasonality and weed management practices effects on the functional structure of weed communities in French vineyards"

**Supplementary table 1** Climate and soil characteristics for each wine-growing region. Climate characteristics data have been extracted from the WorldClim database (Hijmans et al., 2005). Soil characteristics data have been extracted from the Soilgrids dataset at 250 m resolution (Hengl et al., 2017) based on the locations of vineyard plots.

| Climate and soil variables | **Champagne** | **Languedoc** | **Rhône** |
| --- | --- | --- | --- |
| **Climate** | | | |
| Mean annual temperature (°C) | 10.1 | 14.1 | 11.4 |
| Annual rainfall (mm) | 657 | 686 | 776 |
| **Soil** | | | |
| Soil organic carbon content (%) | 16.5 +/- 2.5 | 16.4 +/- 1.3 | 19.7 +/- 1.9 |
| pH index measured in water solutions PHE | 7.1 +/- 0.1 | 7.5 +/- 0.1 | 6.7 +/- 0.3 |
| Cation-exchange capacity (CEC) | 17.4 +/- 1 | 16.1 +/- 1.4 | 17 +/- 2.1 |
| Bulk density (fine heart) (kg/m^3^) TFI | 1387.3 +/- 21.7 | 1527.6 +/- 14.7 | 1473 +/- 28.2 |
| Volumetric percentage of coarse fragments (> 2 mm) (%) PGR | 13.2 +/- 1.2 | 12.5 +/- 2.1 | 13 +/- 1.2 |
| Weight percentage of sand particles (0.05 – 2 mm) (%) PSA | 30.4 +/- 3.8 | 33.7 +/- 2 | 36.7 +/- 3.7 |
| Weight percentage of silt particles (2.10^-4^ – 0.05 mm) (%) PL | 45.7 +/- 3.6 | 38.9 +/- 1.2 | 37.7 +/- 2.2 |
| Weight percentage of clay particles (< 2.10^-4^ mm) (%) PAR | 23.6 +/- 1.8 | 27.4 +/- 2.1 | 25.5 +/- 2 |

|  | **Contribution to axis 1 (%)** | **Contribution to axis 2 (%)** |
| --- | --- | --- |
| **Till.R** | 75 | 1 |
| **Chem.R** | 64 | 4 |
| **Till.IR** | 48 | 19 |
| **Chem.IR** | 18 | 42 |
| **Mow.IR** | 7 | 87 |
| **Years** | 4 | 5 |
| **D** | 3 | 2 |

**Supplementary table 2** Contributions of explaining variables to the PCA of weed management practices and temporal variables (Figure 4). D, number of days since the 1^st^ January of the same year

**Supplementary table 3** Means and standard deviations of community weighted means for each region. Cha, Champagne ; Lan, Languedoc ; Rhô, Rhône ; Sd, standard deviation.

| **Region** | **SLA CWM (mm²/mg)** | | **Maximum height CWM (m)** | | **Seed mass CWM (g)** | | **Flowering onset CWM (months)** | | **Flowering duration CWM (months)** | | **Lateral spread CWM** | |
| --- | --- | --- | --- | --- | --- | --- | --- | --- | --- | --- | --- | --- |
|  | **Mean** | **Sd** | **Mean** | **Sd** | **Mean** | **Sd** | **Mean** | **Sd** | **Mean** | **Sd** | **Mean** | **Sd** |
| **Cha** | 30,0 | 3,1 | 0,73 | 0,16 | 1,410^-03^ | 7,010^-04^ | 3,3 | 0,9 | 8,1 | 1,4 | 1,63 | 0,47 |
| **Lan** | 24,4 | 2,9 | 0,98 | 0,24 | 3,210^-03^ | 3,910^-03^ | 4,8 | 0,9 | 4,5 | 1,0 | 1,81 | 0,90 |
| **Rhô** | 26,7 | 3,6 | 0,80 | 0,20 | 1,810^-03^ | 1,810^-03^ | 4,1 | 1,1 | 6,3 | 1,7 | 1,65 | 0,58 |

**Supplementary table 4** Means and standard deviations of community weighted variance for each region. Cha, Champagne ; Lan, Languedoc ; Rhô, Rhône ; Sd, standard deviation

| **Region** | **SLA CWV (mm²/mg)** | | **Maximum height CWV (m)** | | **Seed mass CWV (g)** | | **Flowering onset CWV (months)** | | **Flowering duration CWV (months)** | | **Lateral spread CWV** | |
| --- | --- | --- | --- | --- | --- | --- | --- | --- | --- | --- | --- | --- |
|  | **Mean** | **Sd** | **Mean** | **Sd** | **Mean** | **Sd** | **Mean** | **Sd** | **Mean** | **Sd** | **Mean** | **Sd** |
| **Cha** | 43,0 | 27,8 | 0,15 | 0,11 | 9,410^-06^ | 1,310^-05^ | 3,5 | 1,9 | 9,5 | 4,3 | 0,88 | 0,67 |
| **Lan** | 23,3 | 17,7 | 0,15 | 0,09 | 2,910^-04^ | 8,610^-04^ | 1,6 | 0,9 | 3,9 | 3,9 | 1,47 | 1,25 |
| **Rhô** | 33,9 | 22,0 | 0,13 | 0,08 | 1,310^-05^ | 2,810^-05^ | 2,8 | 1,7 | 7,0 | 4,2 | 1,24 | 1,02 |

**Supplementary figure 1** Community Weighted Means (CWM) along the first two axes of the weed management practices and temporal variables PCA (Figure 4) in Languedoc. The SLA CWM, maximum height CWM, seed mass CWM of weed community are displayed along the first PCA axis (a, c, e) and along the second PCA axis (b, d, f) respectively. The first PCA axis opposed chemical weeding of rows (Chem.R, negative coordinates,) and tillage of rows and inter-rows (Till.IR, Till.R, positive coordinates). The second PCA axis opposed mowing of inter-rows (Mow.IR, negative coordinates) to combination of tillage and chemical weeding of inter-rows (Chem.IR + Till.IR, positive coordinates). Each point is a weed community. The color of each point specify the weed management of inter-rows (C, chemical weeding ; CT, combination of chemical weeding and tillage ; M, mowing ; T, Tillage). The shape indicate the weed management of rows (C, chemical weeding ; CT, combination of chemical weeding and tillage ; T, Tillage).

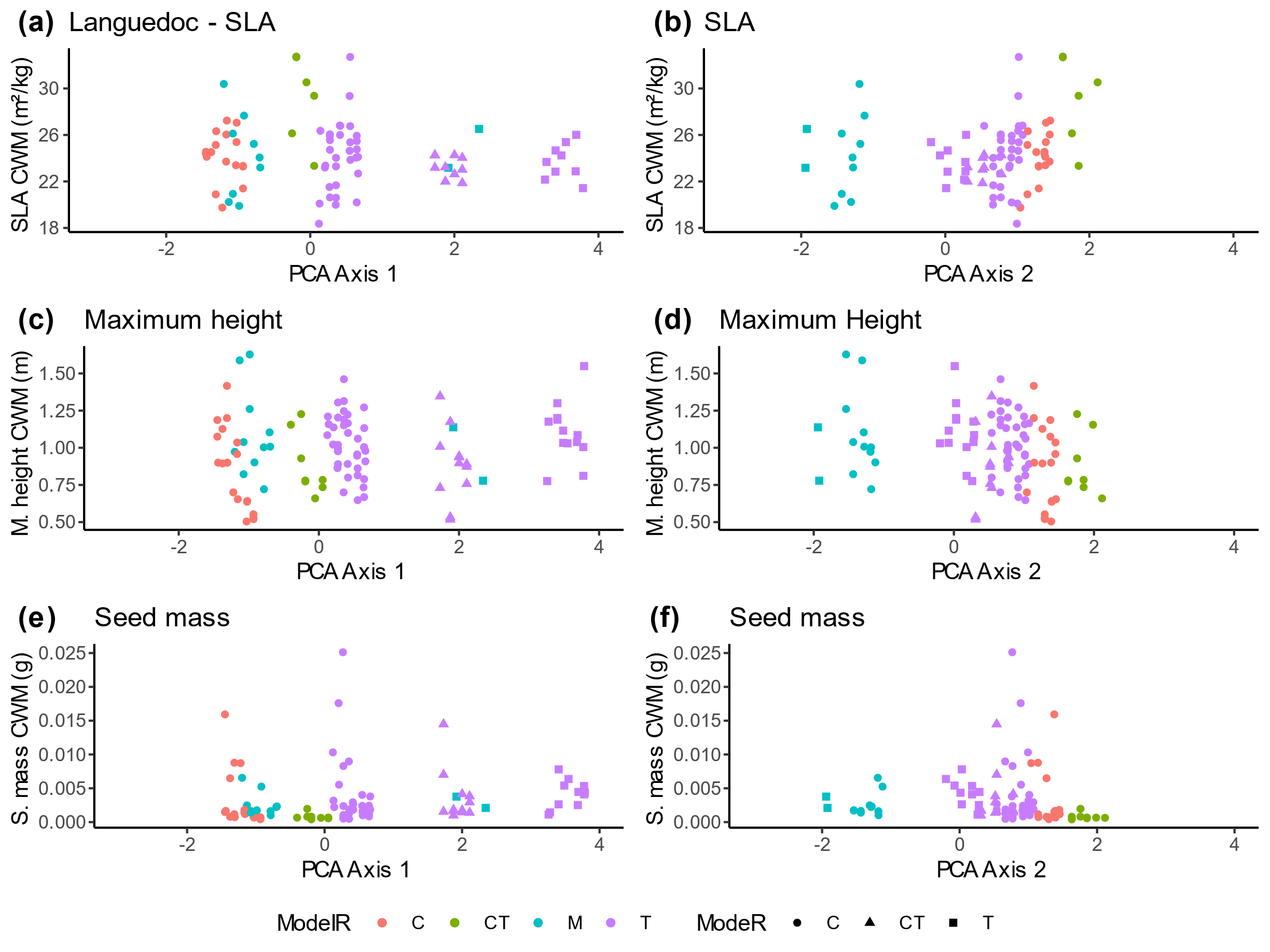

**Supplementary figure 2** Community Weighted Means (CWM) along the first two axes of the weed management practices and temporal variables PCA (Figure 4) in Languedoc. The lateral spread CWM, flowering onset CWM, flowering duration CWM of weed community are displayed along the first PCA axis (a, c, e) and along the second PCA axis (b, d, f) respectively. The first PCA axis opposed chemical weeding of rows (Chem.R, negative coordinates,) and tillage of rows and inter-rows (Till.IR, Till.R, positive coordinates). The second PCA axis opposed mowing of inter-rows (Mow.IR, negative coordinates) to combination of tillage and chemical weeding of inter-rows (Chem.IR + Till.IR, positive coordinates). Each point is a weed community. The color of each point specify the weed management of inter-rows (C, chemical weeding ; CT, combination of chemical weeding and tillage ; M, mowing ; T, Tillage). The shape indicate the weed management of rows (C, chemical weeding ; CT, combination of chemical weeding and tillage ; T, Tillage).

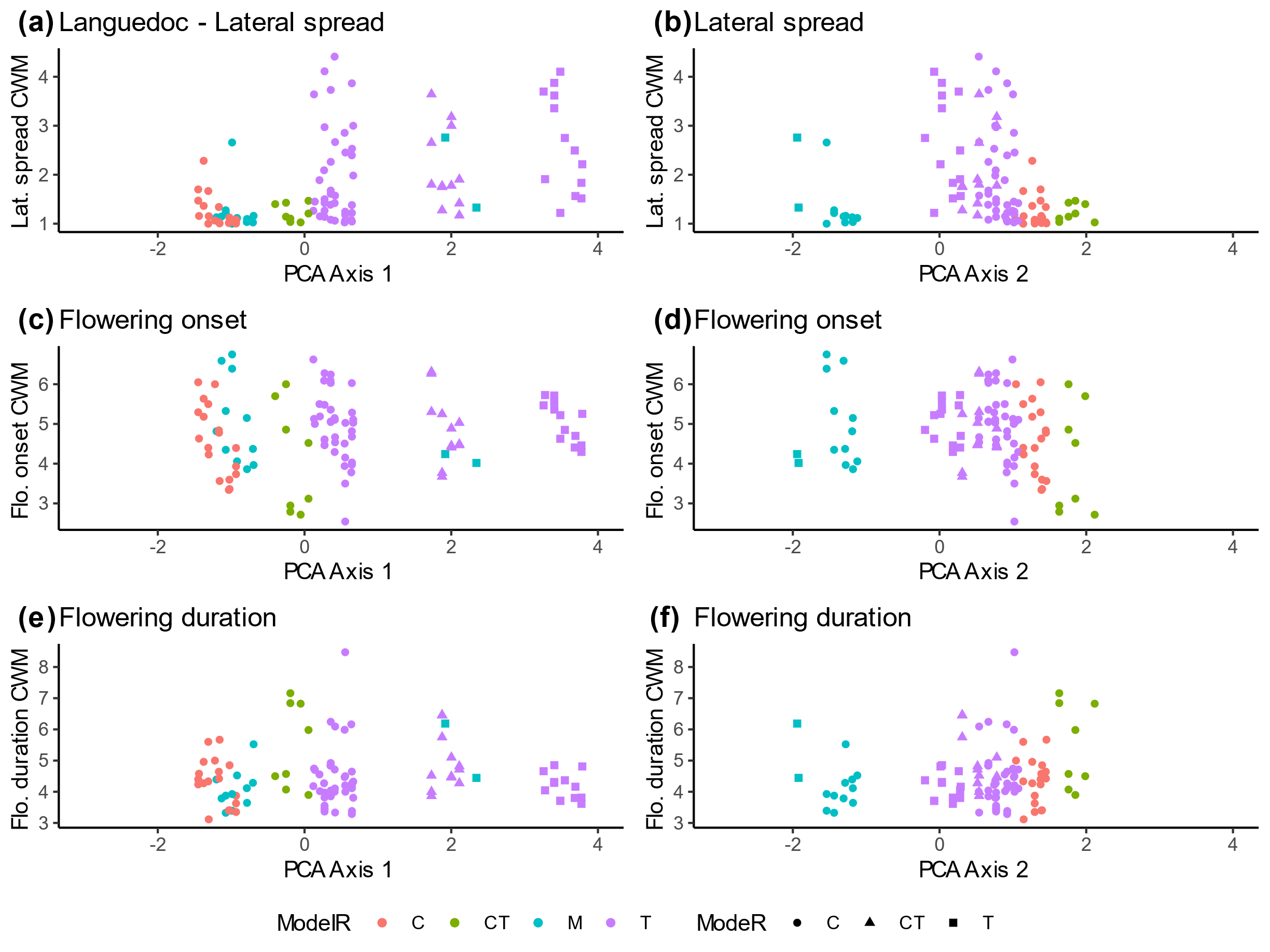

**Supplementary figure 3** Community Weighted Means (CWM) along the first two axes of the weed management practices and temporal variables PCA (Figure 4) in Champagne. The SLA CWM, maximum height CWM, seed mass CWM of weed community are displayed along the first PCA axis (a, c, e) and along the second PCA axis (b, d, f) respectively. The first PCA axis opposed chemical weeding of rows (Chem.R, negative coordinates,) and tillage of rows and inter-rows (Till.IR, Till.R, positive coordinates). The second PCA axis opposed mowing of inter-rows (Mow.IR, negative coordinates) to combination of tillage and chemical weeding of inter-rows (Chem.IR + Till.IR, positive coordinates). Each point is a weed community. The color of each point specify the weed management of inter-rows (C, chemical weeding ; CT, combination of chemical weeding and tillage ; CM, combination of chemical weeding and mowing ; TM, combination of tillage and mowing ; M, mowing ; T, Tillage). The shape indicate the weed management of rows (C, chemical weeding ; CT, combination of chemical weeding and tillage ; T, Tillage).

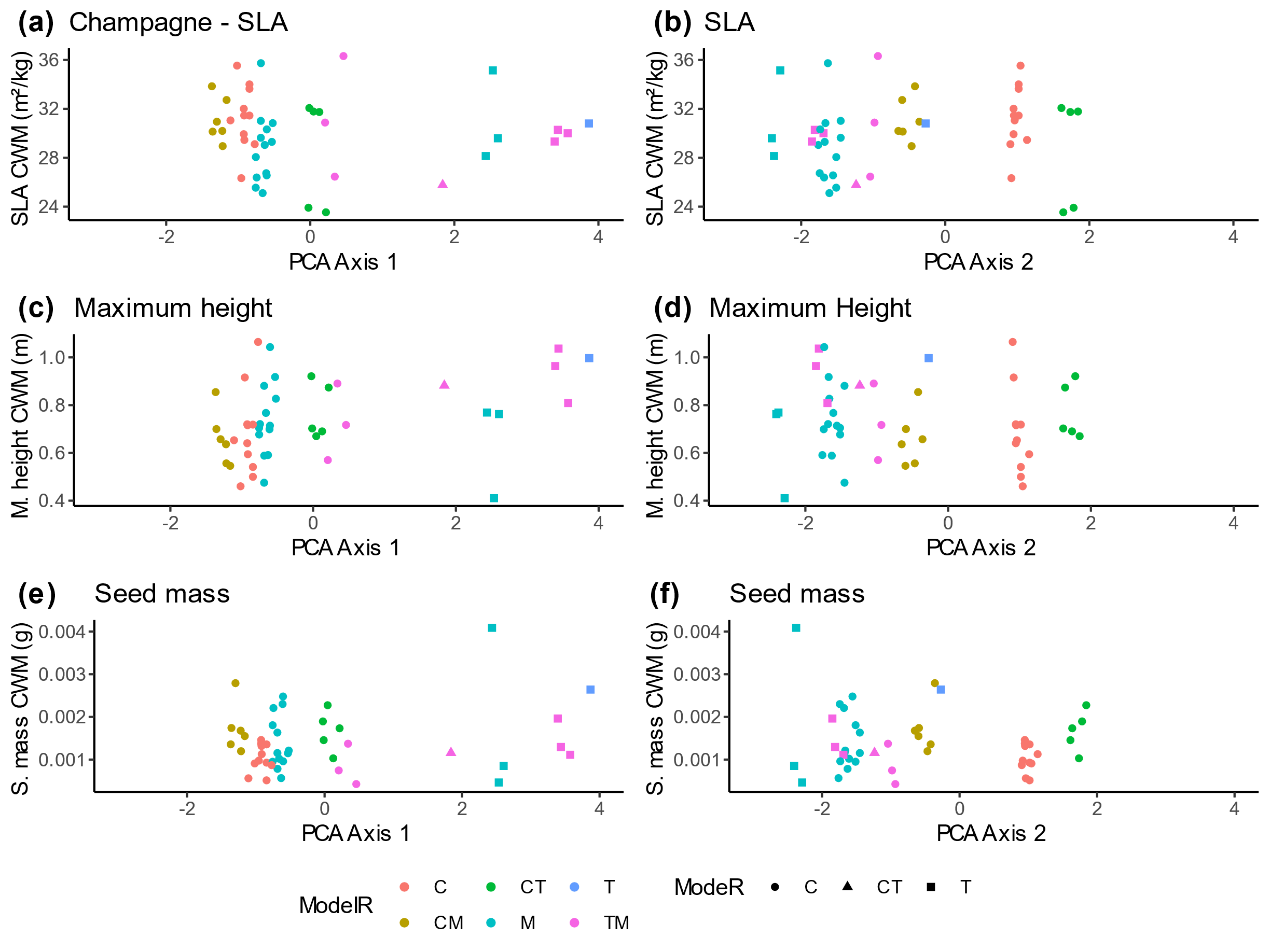

**Supplementary figure 4** Community Weighted Means (CWM) along the first two axes of the weed management practices and temporal variables PCA (Figure 4) in Champagne. The lateral spread CWM, flowering onset CWM, flowering duration CWM of weed community are displayed along the first PCA axis (a, c, e) and along the second PCA axis (b, d, f) respectively. The first PCA axis opposed chemical weeding of rows (Chem.R, negative coordinates,) and tillage of rows and inter-rows (Till.IR, Till.R, positive coordinates). The second PCA axis opposed mowing of inter-rows (Mow.IR, negative coordinates) to combination of tillage and chemical weeding of inter-rows (Chem.IR + Till.IR, positive coordinates). Each point is a weed community. The color of each point specify the weed management of inter-rows (C, chemical weeding ; CT, combination of chemical weeding and tillage ; CM, combination of chemical weeding and mowing ; TM, combination of tillage and mowing ; M, mowing ; T, Tillage). The shape indicate the weed management of rows (C, chemical weeding ; CT, combination of chemical weeding and tillage ; T, Tillage).

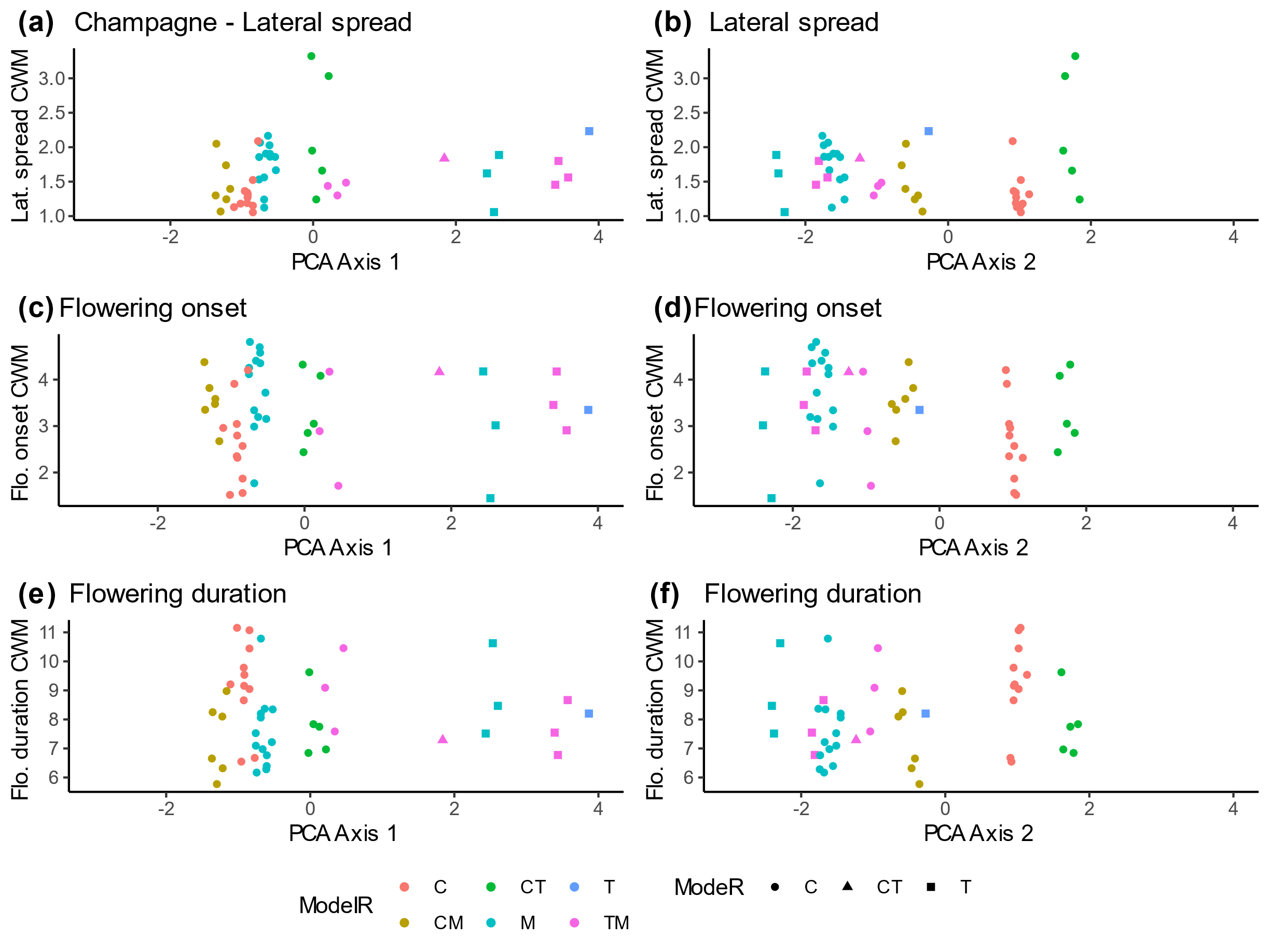

**Supplementary figure 5** Community Weighted Means (CWM) along the first two axes of the weed management practices and temporal variables PCA (Figure 4) in Rhône. The SLA CWM, maximum height CWM, seed mass CWM of weed community are displayed along the first PCA axis (a, c, e) and along the second PCA axis (b, d, f) respectively. The first PCA axis opposed chemical weeding of rows (Chem.R, negative coordinates,) and tillage of rows and inter-rows (Till.IR, Till.R, positive coordinates). The second PCA axis opposed mowing of inter-rows (Mow.IR, negative coordinates) to combination of tillage and chemical weeding of inter-rows (Chem.IR + Till.IR, positive coordinates). Each point is a weed community. The color of each point specify the weed management of inter-rows (C, chemical weeding ; CT, combination of chemical weeding and tillage ; CM, combination of chemical weeding and mowing ; M, mowing ; T, Tillage). The shape indicate the weed management of rows (C, chemical weeding ; CT, combination of chemical weeding and tillage ; T, Tillage).

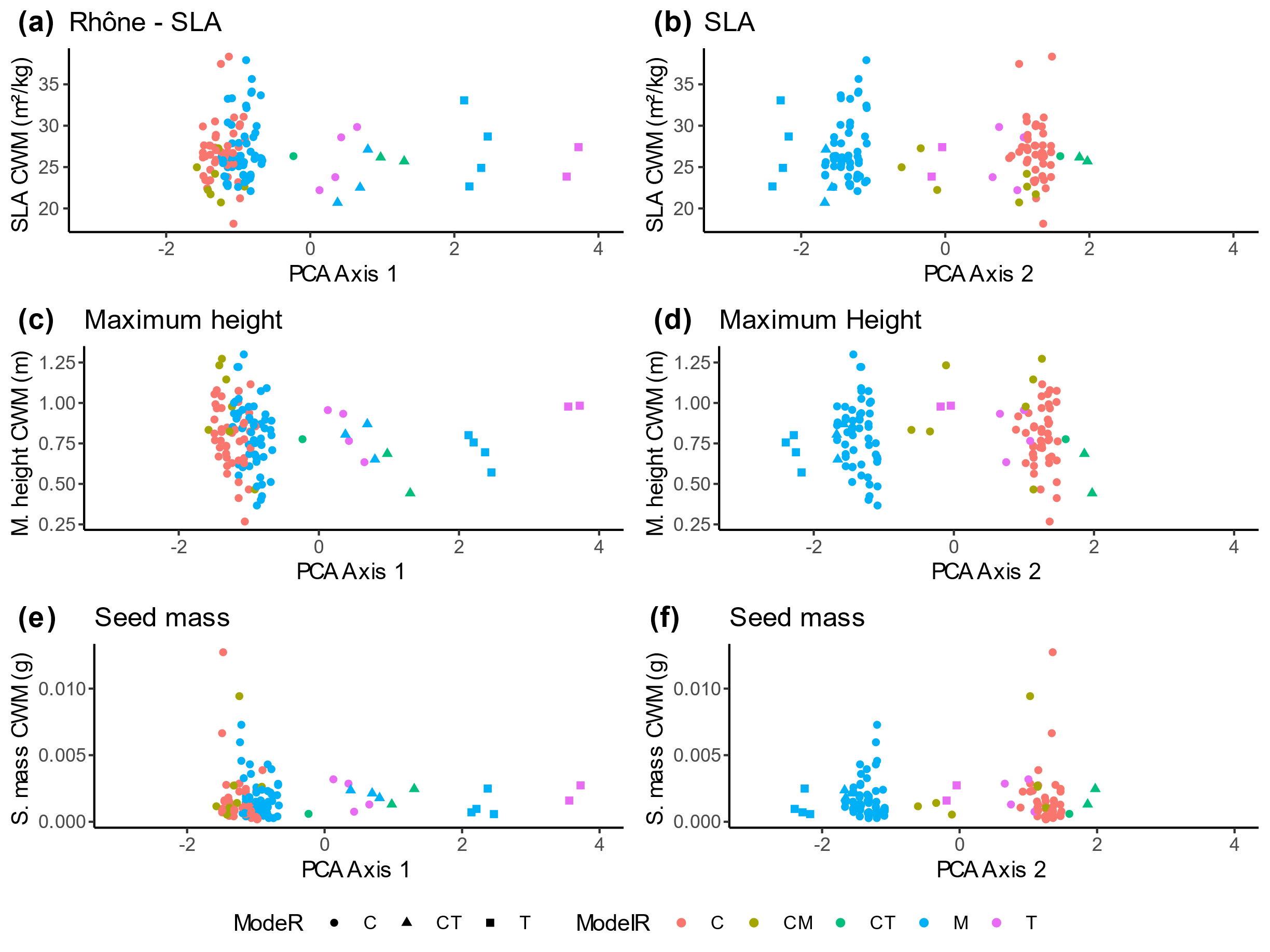

**Supplementary figure 4** Community Weighted Means (CWM) along the first two axes of the weed management practices and temporal variables PCA (Figure 4) in Rhône. The lateral spread CWM, flowering onset CWM, flowering duration CWM of weed community are displayed along the first PCA axis (a, c, e) and along the second PCA axis (b, d, f) respectively. The first PCA axis opposed chemical weeding of rows (Chem.R, negative coordinates,) and tillage of rows and inter-rows (Till.IR, Till.R, positive coordinates). The second PCA axis opposed mowing of inter-rows (Mow.IR, negative coordinates) to combination of tillage and chemical weeding of inter-rows (Chem.IR + Till.IR, positive coordinates). Each point is a weed community. The color of each point specify the weed management of inter-rows (C, chemical weeding ; CT, combination of chemical weeding and tillage ; CM, combination of chemical weeding and mowing ; M, mowing ; T, Tillage). The shape indicate the weed management of rows (C, chemical weeding ; CT, combination of chemical weeding and tillage ; T, Tillage).

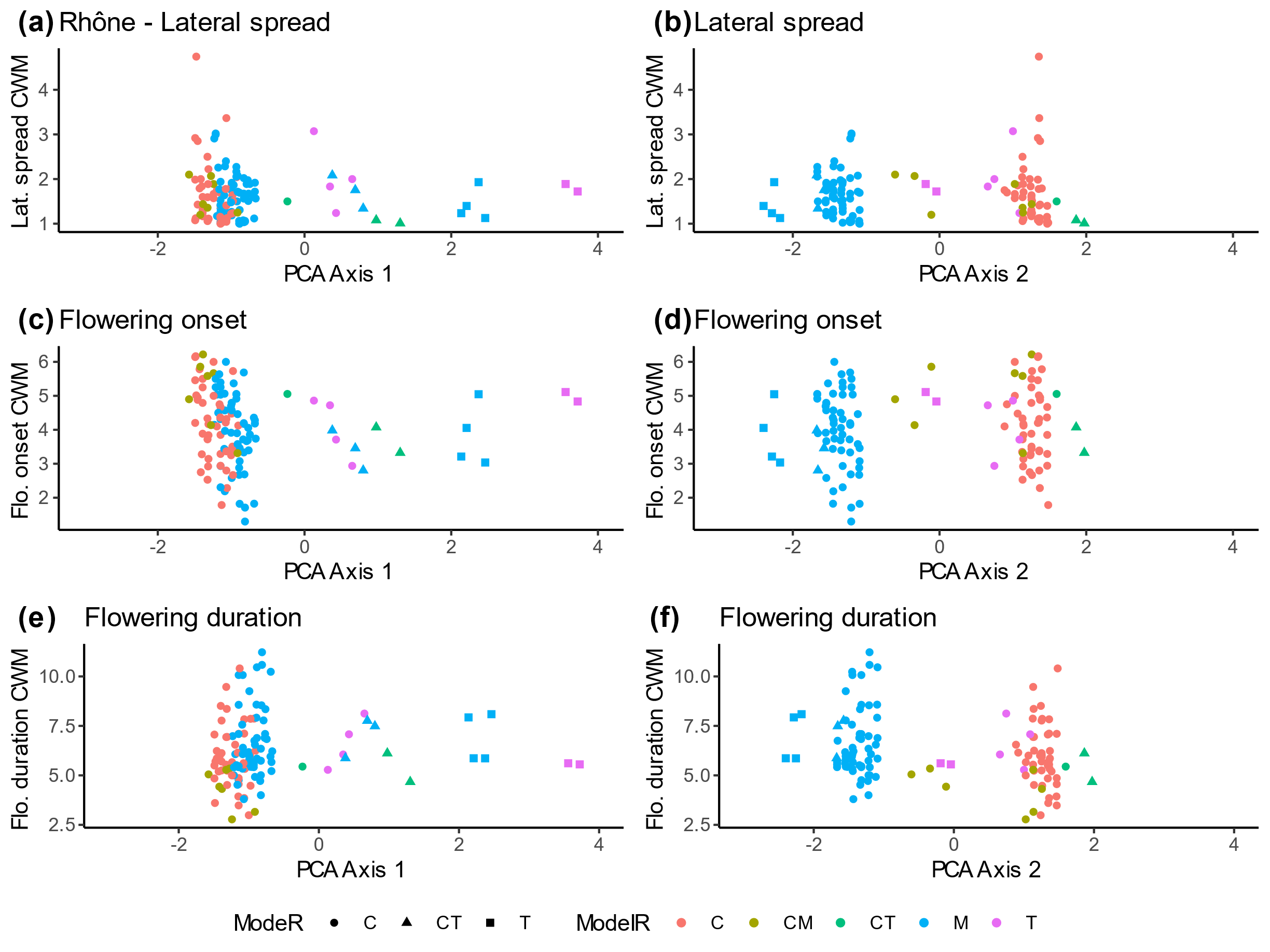

**Supplementary figure 7** Effect sizes of Community Weighted Variance (CWV) along the first two axes of the weed management practices and temporal variables PCA (Figure 4) in Languedoc. The effect size of SLA CWV, maximum height CWV and seed mass CWV of weed community are displayed along the first PCA axis (a, c, e) and along the second PCA axis (b, d, f) respectively. The first PCA axis opposed chemical weeding of rows (Chem.R, negative coordinates,) and tillage of rows and inter-rows (Till.IR, Till.R, positive coordinates). The second PCA axis opposed mowing of inter-rows (Mow.IR, negative coordinates) to combination of tillage and chemical weeding of inter-rows (Chem.IR + Till.IR, positive coordinates). Each point is a weed community. The color of each point specify the weed management of inter-rows (C, chemical weeding ; CT, combination of chemical weeding and tillage ; M, mowing ; T, Tillage). The shape indicate the weed management of rows (C, chemical weeding ; CT, combination of chemical weeding and tillage ; T, Tillage). The results of the two-tailed Wilcoxon signed-ranks test (W) is specified. High and low ES values quantify respectively divergent and convergent functional structure of weed communities. Significance of Wilcoxon tests are referred as following: * *p* < 0.05 ; ** *p* < 0.01 ; *** *p* < 0.001, ns ; non significant.

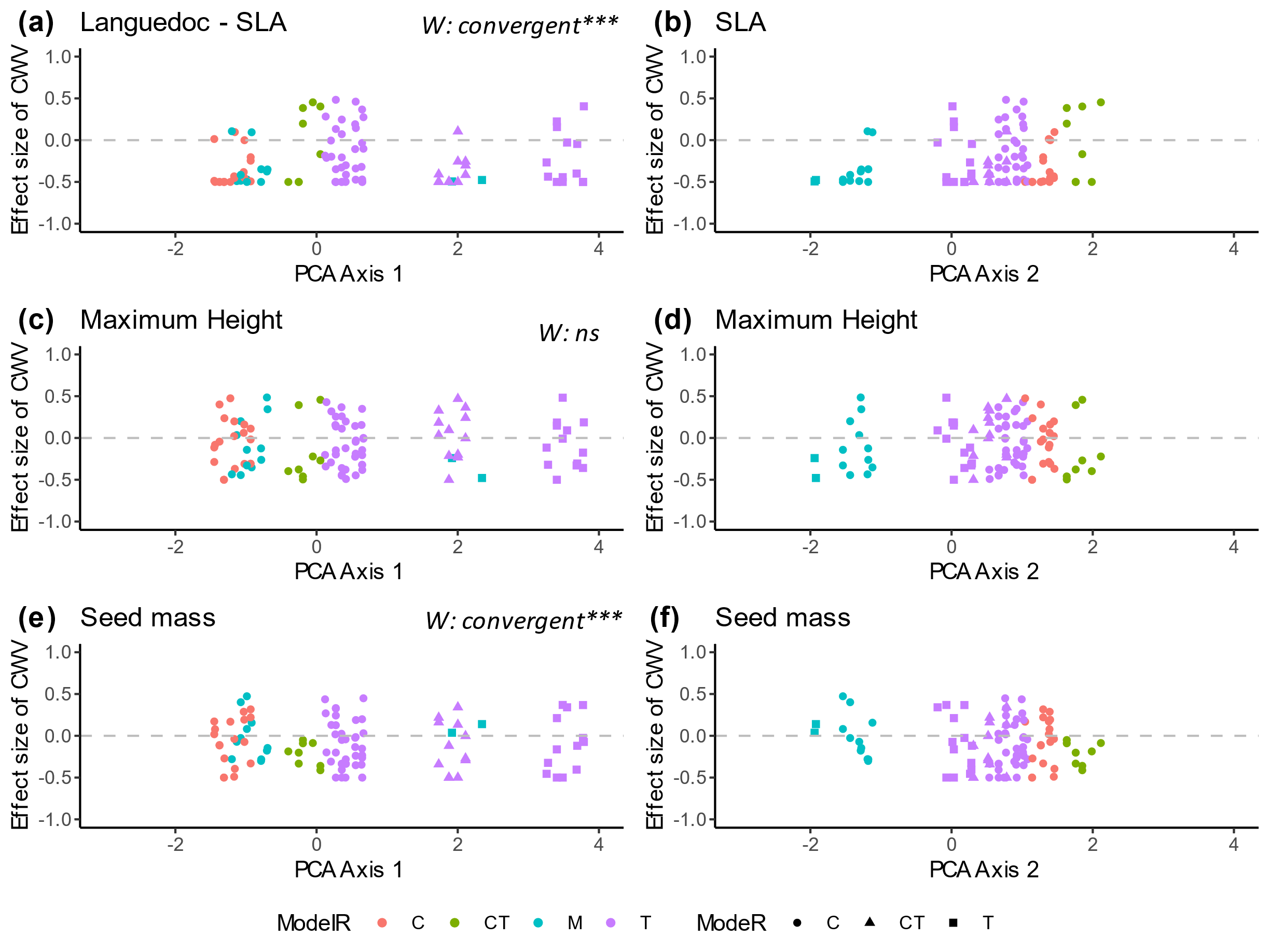

**Supplementary figure 8** Effect sizes of Community Weighted Variance (CWV) along the first two axes of the weed management practices and temporal variables PCA (Figure 4) in Languedoc. The effect size of lateral spread CWV, flowering onset CWV and flowering duration CWV of weed community are displayed along the first PCA axis (a, c, e) and along the second PCA axis (b, d, f) respectively. The first PCA axis opposed chemical weeding of rows (Chem.R, negative coordinates,) and tillage of rows and inter-rows (Till.IR, Till.R, positive coordinates). The second PCA axis opposed mowing of inter-rows (Mow.IR, negative coordinates) to combination of tillage and chemical weeding of inter-rows (Chem.IR + Till.IR, positive coordinates). Each point is a weed community. The color of each point specify the weed management of inter-rows (C, chemical weeding ; CT, combination of chemical weeding and tillage ; M, mowing ; T, Tillage). The shape indicate the weed management of rows (C, chemical weeding ; CT, combination of chemical weeding and tillage ; T, Tillage). The results of the two-tailed Wilcoxon signed-ranks test (W) is specified. High and low ES values quantify respectively divergent and convergent functional structure of weed communities. Significance of Wilcoxon tests are referred as following: * *p* < 0.05 ; ** *p* < 0.01 ; *** *p* < 0.001, ns ; non significant.

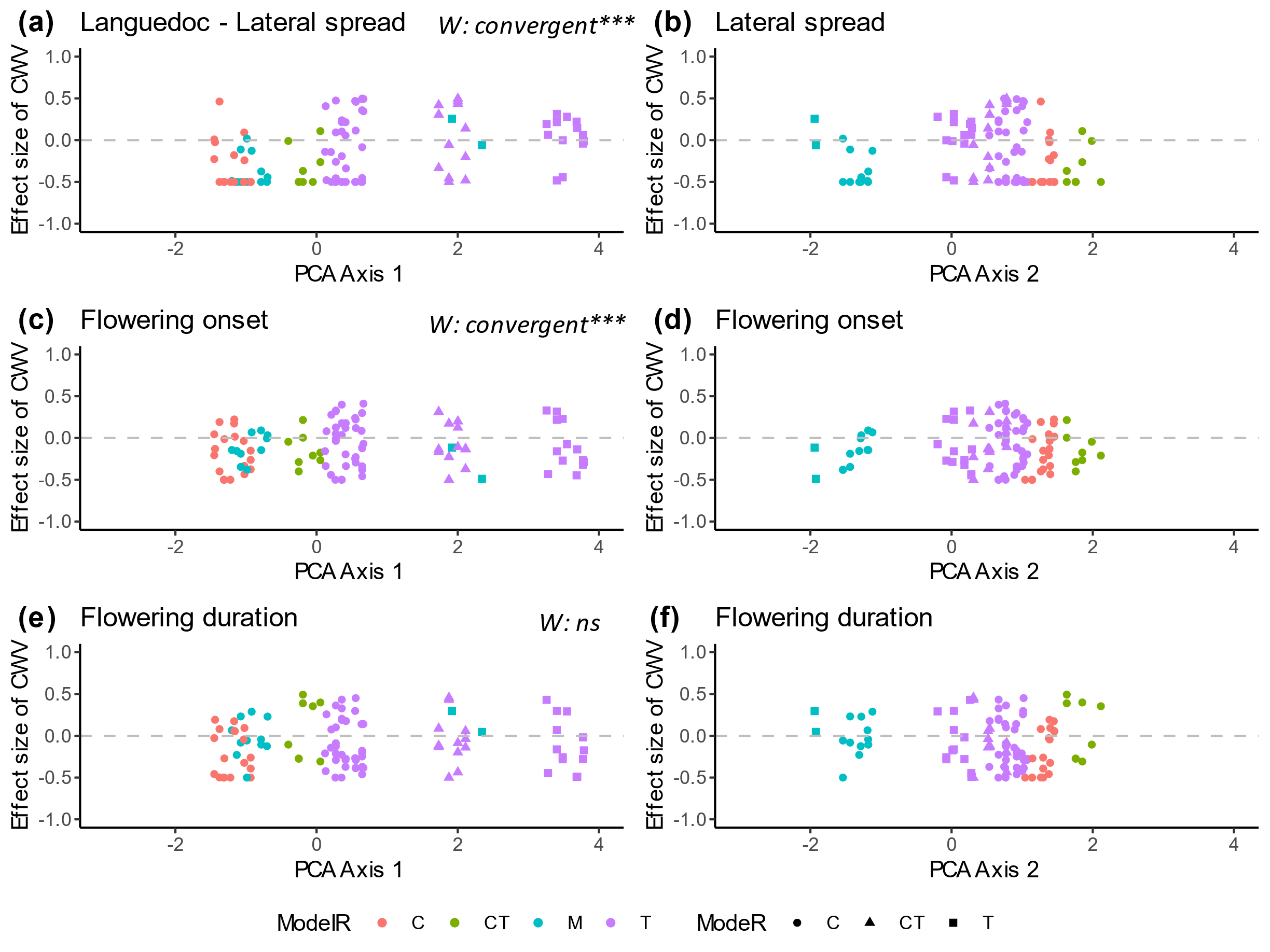

**Supplementary figure 9** Effect sizes of Community Weighted Variance (CWV) along the first two axes of the weed management practices and temporal variables PCA (Figure 4) in Champagne. The effect size of SLA CWV, maximum height CWV and seed mass CWV of weed community are displayed along the first PCA axis (a, c, e) and along the second PCA axis (b, d, f) respectively. The first PCA axis opposed chemical weeding of rows (Chem.R, negative coordinates,) and tillage of rows and inter-rows (Till.IR, Till.R, positive coordinates). The second PCA axis opposed mowing of inter-rows (Mow.IR, negative coordinates) to combination of tillage and chemical weeding of inter-rows (Chem.IR + Till.IR, positive coordinates). Each point is a weed community. The color of each point specify the weed management of inter-rows (C, chemical weeding ; CT, combination of chemical weeding and tillage ; M, mowing ; T, Tillage). The color of each point specify the weed management of inter-rows (C, chemical weeding ; CT, combination of chemical weeding and tillage ; CM, combination of chemical weeding and mowing ; TM, combination of tillage and mowing ; M, mowing ; T, Tillage). The results of the two-tailed Wilcoxon signed-ranks test (W) is specified. High and low ES values quantify respectively divergent and convergent functional structure of weed communities. Significance of Wilcoxon tests are referred as following: * *p* < 0.05 ; ** *p* < 0.01 ; *** *p* < 0.001, ns ; non significant.

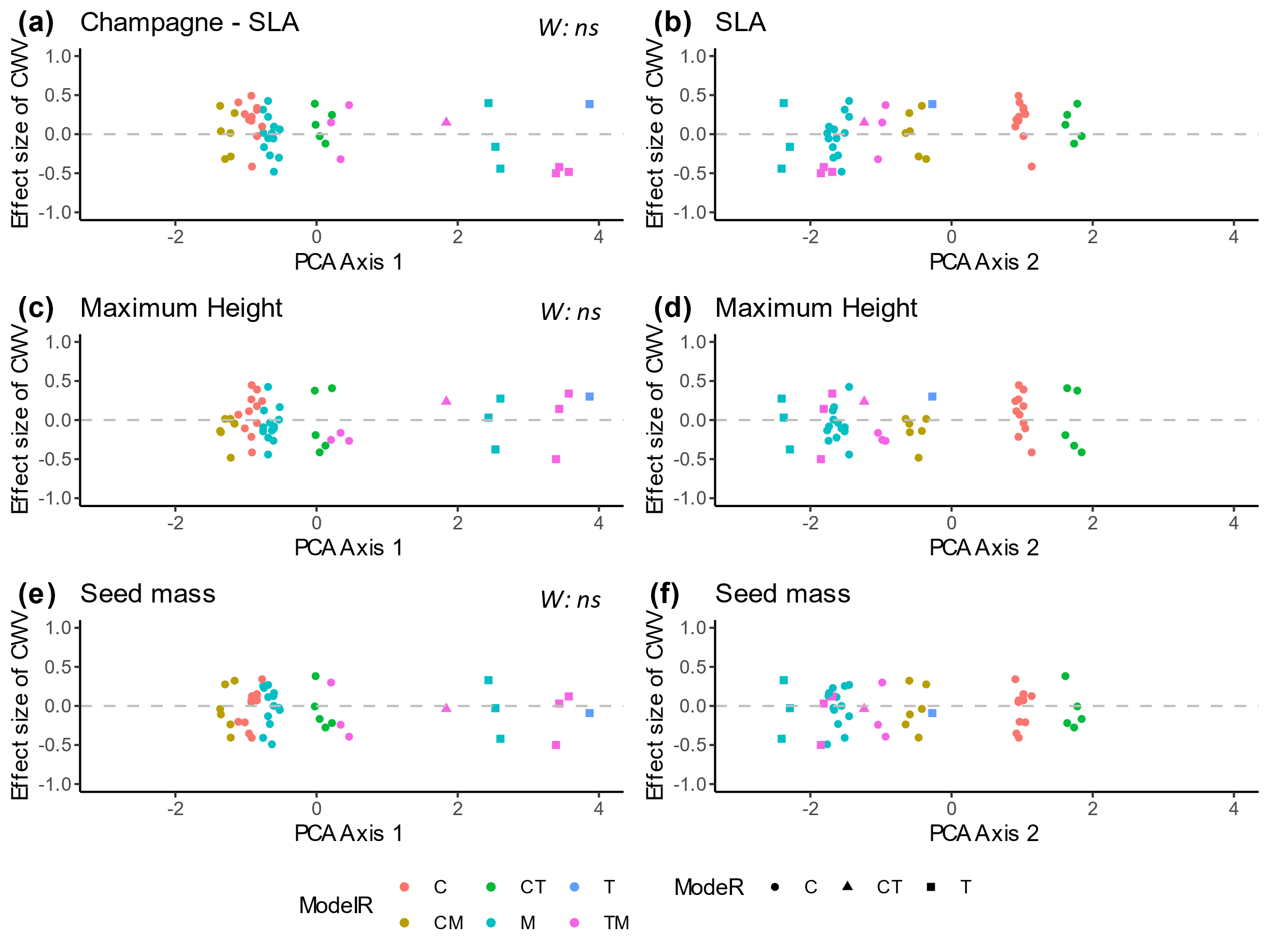

**Supplementary figure 10** Effect sizes of Community Weighted Variance (CWV) along the first two axes of the weed management practices and temporal variables PCA (Figure 4) in Champagne. The effect size of lateral spread CWV, flowering onset CWV and flowering duration CWV of weed community are displayed along the first PCA axis (a, c, e) and along the second PCA axis (b, d, f) respectively. The first PCA axis opposed chemical weeding of rows (Chem.R, negative coordinates,) and tillage of rows and inter-rows (Till.IR, Till.R, positive coordinates). The second PCA axis opposed mowing of inter-rows (Mow.IR, negative coordinates) to combination of tillage and chemical weeding of inter-rows (Chem.IR + Till.IR, positive coordinates). Each point is a weed community. The color of each point specify the weed management of inter-rows (C, chemical weeding ; CT, combination of chemical weeding and tillage ; CM, combination of chemical weeding and mowing ; TM, combination of tillage and mowing ; M, mowing ; T, Tillage). The shape indicate the weed management of rows (C, chemical weeding ; CT, combination of chemical weeding and tillage ; T, Tillage). The results of the two-tailed Wilcoxon signed-ranks test (W) is specified. High and low ES values quantify respectively divergent and convergent functional structure of weed communities. Significance of Wilcoxon tests are referred as following: * *p* < 0.05 ; ** *p* < 0.01 ; *** *p* < 0.001, ns ; non significant.

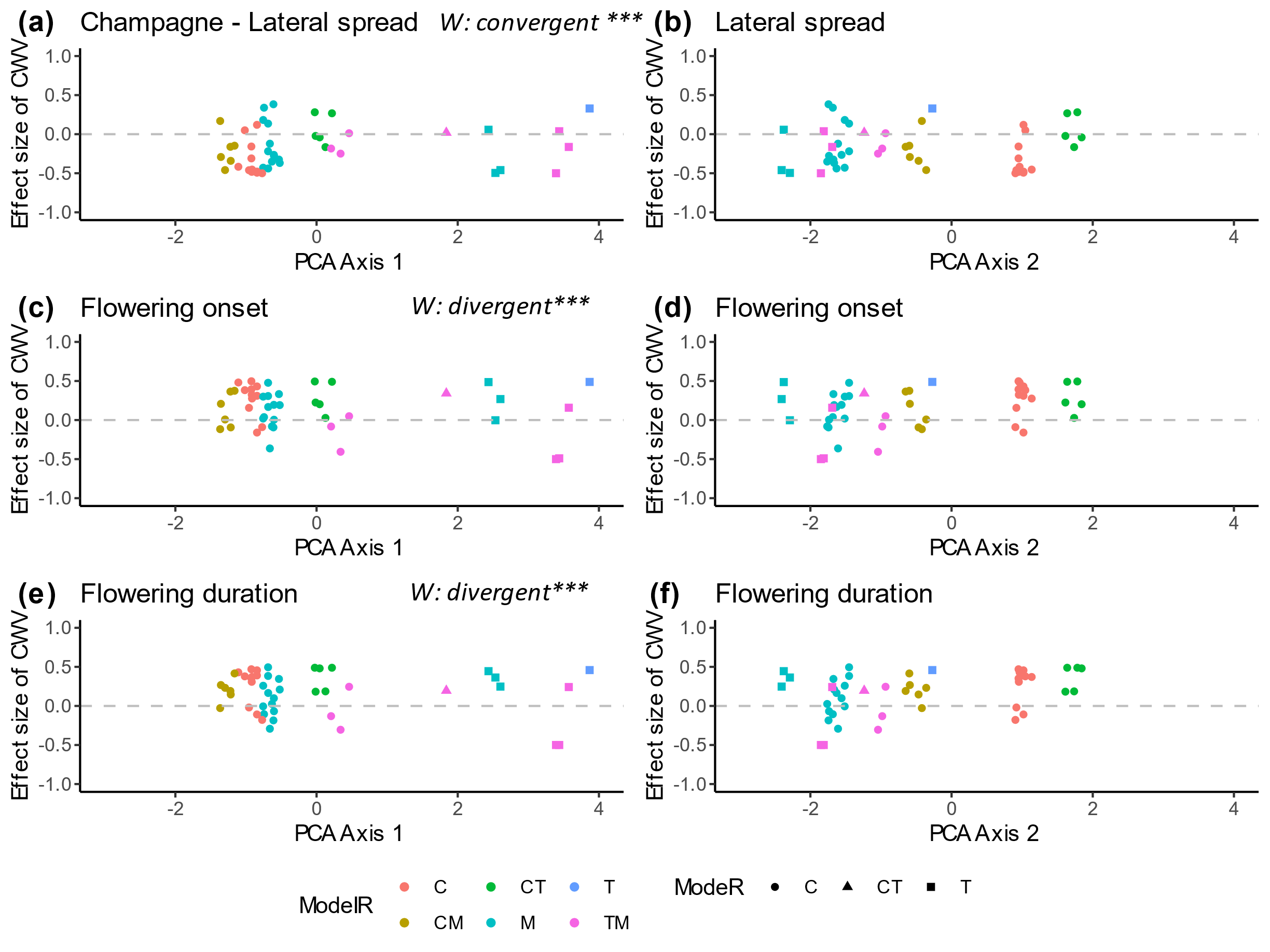

**Supplementary figure 11** Effect sizes of Community Weighted Variance (CWV) along the first two axes of the weed management practices and temporal variables PCA (Figure 4) in Rhône. The effect size of SLA CWV, maximum height CWV and seed mass CWV of weed community are displayed along the first PCA axis (a, c, e) and along the second PCA axis (b, d, f) respectively. The first PCA axis opposed chemical weeding of rows (Chem.R, negative coordinates,) and tillage of rows and inter-rows (Till.IR, Till.R, positive coordinates). The second PCA axis opposed mowing of inter-rows (Mow.IR, negative coordinates) to combination of tillage and chemical weeding of inter-rows (Chem.IR + Till.IR, positive coordinates). Each point is a weed community. The color of each point specify the weed management of inter-rows (C, chemical weeding ; CT, combination of chemical weeding and tillage ; M, mowing ; T, Tillage). The color of each point specify the weed management of inter-rows (C, chemical weeding ; CT, combination of chemical weeding and tillage ; CM, combination of chemical weeding and mowing ; M, mowing ; T, Tillage). The results of the two-tailed Wilcoxon signed-ranks test (W) is specified. High and low ES values quantify respectively divergent and convergent functional structure of weed communities. Significance of Wilcoxon tests are referred as following: * *p* < 0.05 ; ** *p* < 0.01 ; *** *p* < 0.001, ns ; non significant.

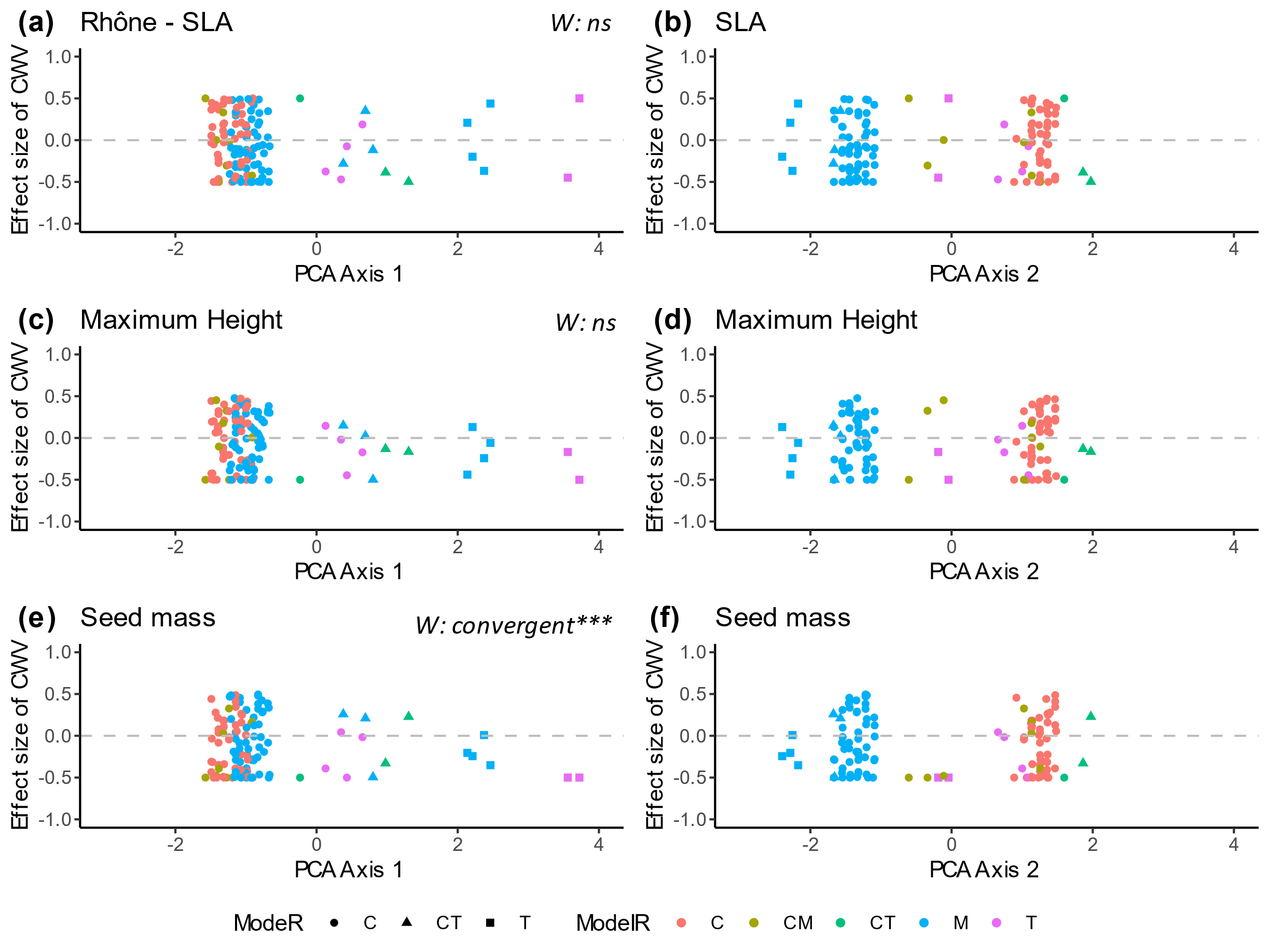

**Supplementary figure 12** Effect sizes of Community Weighted Variance (CWV) along the first two axes of the weed management practices and temporal variables PCA (Figure 4) in Rhône. The effect size of lateral spread CWV, flowering onset CWV and flowering duration CWV of weed community are displayed along the first PCA axis (a, c, e) and along the second PCA axis (b, d, f) respectively. The first PCA axis opposed chemical weeding of rows (Chem.R, negative coordinates,) and tillage of rows and inter-rows (Till.IR, Till.R, positive coordinates). The second PCA axis opposed mowing of inter-rows (Mow.IR, negative coordinates) to combination of tillage and chemical weeding of inter-rows (Chem.IR + Till.IR, positive coordinates). Each point is a weed community. The color of each point specify the weed management of inter-rows (C, chemical weeding ; CT, combination of chemical weeding and tillage ; CM, combination of chemical weeding and mowing ; M, mowing ; T, Tillage). The shape indicate the weed management of rows (C, chemical weeding ; CT, combination of chemical weeding and tillage ; T, Tillage). The results of the two-tailed Wilcoxon signed-ranks test (W) is specified. High and low ES values quantify respectively divergent and convergent functional structure of weed communities. Significance of Wilcoxon tests are referred as following: * *p* < 0.05 ; ** *p* < 0.01 ; *** *p* < 0.001, ns ; non significant.

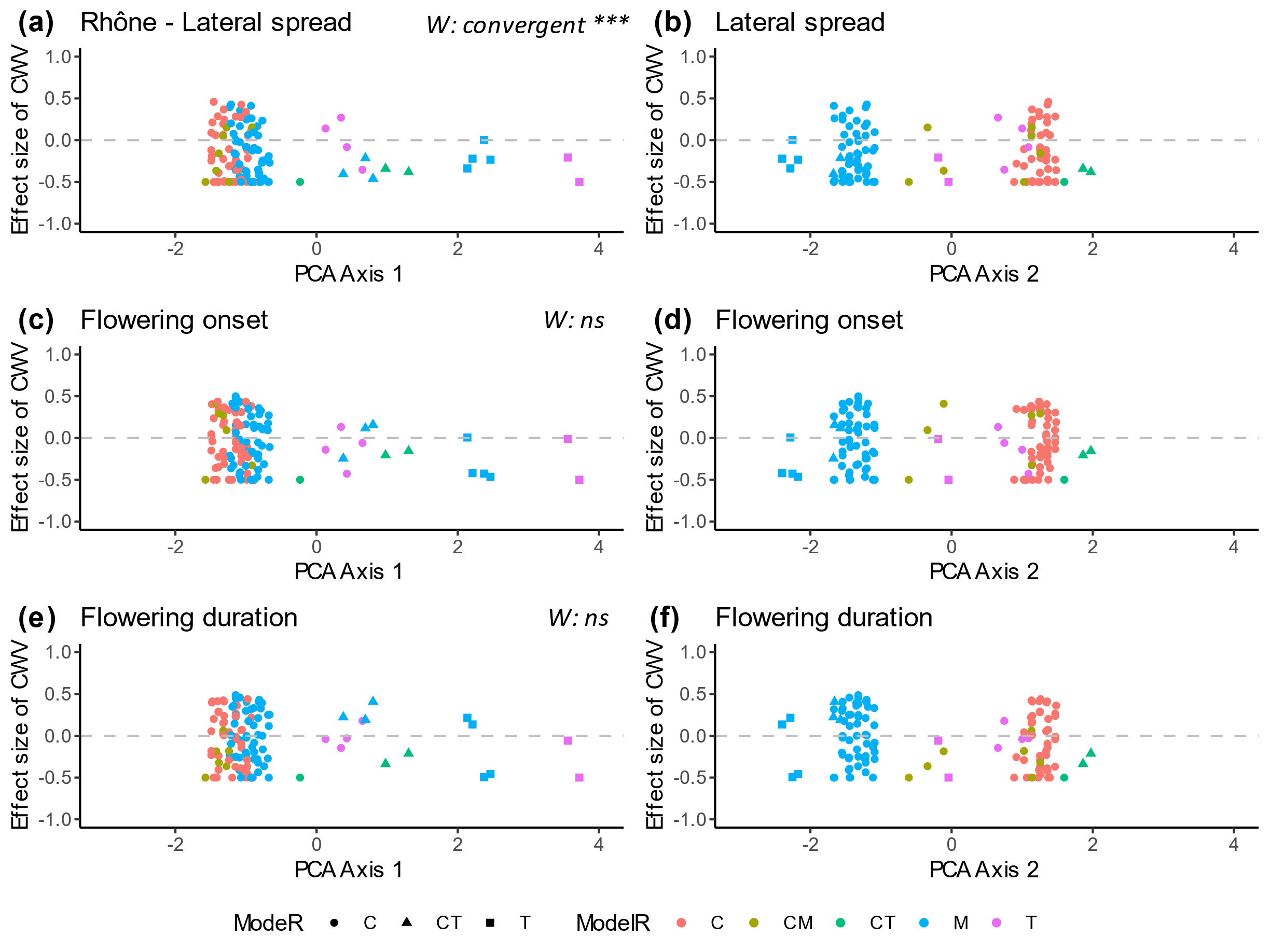

**Supplementary figure 13** Densities of lateral spread community weighted variance in Languedoc from different rows managements (C, chemical weeding ; CT, combination of chemical weeding and tillage ; T, tillage).

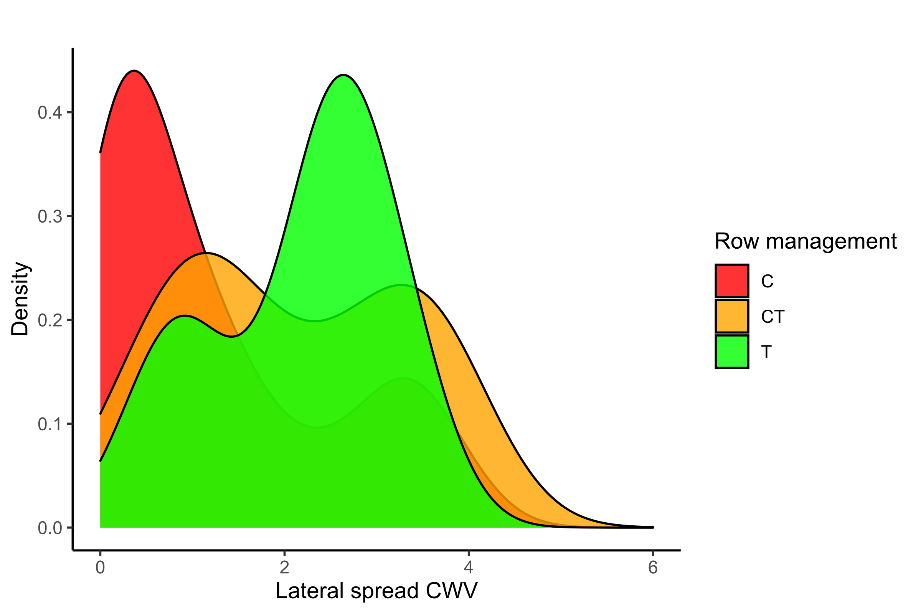
